# Guided maturation of human neuromuscular organoids via electrical stimulation

**DOI:** 10.1101/2025.10.30.685366

**Authors:** Chrysanthi-Maria Moysidou, Inês Afonso Martins, Ismail Amr El-Shimy, Donatella Cea, Christina Bukas, Isra Mekki, Ines Lahmann, Marie Piraud, Enrico Klotzsch, Mina Gouti

## Abstract

Organoids derived from human pluripotent stem cells (hPSCs) are emerging as powerful models for studying development and disease. Despite their physiological relevance, the predictive power of organoids remains limited by the immature state of the constituent cells, posing a major challenge for mechanistic studies of adult physiology and late-onset diseases and disorders. Here, we establish a strategy for enhancing the maturation status of human neuromuscular organoids (NMOs) through chronic Electrical Pulse Stimulation (EPS). We demonstrate that low-frequency EPS, applied early on during NMO development and maintained over several weeks, promotes structural and functional maturation of neuromuscular junctions (NMJs). Independent of stimulation waveform dynamics, EPS-trained NMOs (EPS-NMOs) displayed stronger and more frequent spontaneous contractions that persisted long after stimulation ceased. Quantitative imaging and transcriptomic analyses revealed a robust improvement in EPS-NMO skeletal muscle and neural tissue morphology, coordinated regulation of lineage-specific biomarkers, and upregulation of gene programmes associated with mature neuromuscular function. Mechanobiological measurements further demonstrated increased EPS-NMO tissue stiffness and faster relaxation dynamics, consistent with advanced excitation–contraction coupling and force generation. Collectively, these findings establish EPS as a powerful, non-invasive, and on-demand modality for driving the morphological and functional maturation of complex organoid systems.

## Introduction

Advances in stem cell technology have enabled the generation of three-dimensional (3D) self-organising cellular ensembles, namely organoids.^1,2^ Unlike two-dimensional (2D) cell culture models, human organoids capture the 3D cytoarchitecture, cellular diversity, and intercellular interactions of their in vivo counterparts, thus providing powerful in vitro systems to study human development and disease.^1,3^ As such, organoids are considered paradigm-shifting tools in biomedical research and are increasingly employed in modelling various human tissues, including intestine,^4^ liver,^5,6^ lung,^7,8^ heart,^9^ and brain^10,11^, facilitating in vitro studies of organogenesis and disease at an unprecedented level.^1,12^ Despite these advances, achieving adult-like tissue maturation states in organoids remains a challenge.^2,13,14^ Attaining such levels of maturation is critical for employing organoids in adult-onset disease modelling (e.g., Amyotrophic Lateral Sclerosis (ALS), Alzheimer’s disease (AS), Age-related Macular Degeneration (AMD), cancer), as well as in drug discovery pipelines and personalised medicine applications.^2,15^

Several strategies have been developed to promote organoid maturation, including long term culture to approximate developmental timelines,^3^ co-culture or fusion with other cell types and/or organoids,^16–18^ and transplantation into host tissues.^19–22^ Despite the valuable insight these approaches offer, organoids remain largely constrained to foetal or early post-natal stages,^2,12,23–25^ indicating that essential cues are missing. Recent work has made strides in identifying such factors, highlighting the differential effect of distinct stimuli on tissue-specific maturation pathways.^13,17^ Among the most promising approaches are complex media formulations, hormonal stimulation,^13,26,27^ and integration of biophysical cues,^2,14,28,29^ such as mechanical loading and electrical stimulation, which has emerged as one of the most potent drivers for functional tissue maturation.^30–32^

Bioelectric manipulation of tissues can be traced back to Luigi Galvani’s experiments on frog leg muscle twitches, demonstrating that muscle contraction is driven by electrical stimuli, laying the foundation of modern electrophysiology.^33^ We now know that the fundamental mechanism by which motor neuron action potentials are converted into muscle fibre mechanical contraction is excitation-contraction coupling (ECC).^34–36^ In skeletal muscle, ECC starts at the neuromuscular junction (NMJ) upon arrival of motor neuron action potentials, triggering the release of acetylcholine (Ach) from the presynaptic terminal into the synaptic cleft. Binding of Ach to its receptors at the post-synaptic terminal induces depolarisation of the sarcolemma, which is propagated down the muscle fibres. Such rapid changes in the muscle membrane electrical properties cause release of large amounts of Ca^2+^ into the sarcoplasm and structural conformation of myosin and actin proteins that slide along each other, leading to myofibril contractions.^34,36,37^ Electrical Pulse Stimulation (EPS) is commonly used to study and modulate ECC in in vitro models, emulating motor neuron activation of skeletal muscle cultures that are typically quiescent and/or do not exhibit spontaneous contraction.^38–42^ While EPS has been widely applied to 2D myotube cultures to enhance contractility and metabolic function, it is typically introduced only at late differentiation states and in the absence of motor neuron input.^43,44^ As a result, most studies do not address how bioelectrical cues during early developmental stages might influence reciprocal neuron-muscle signalling, self-organization, and NMJ formation and maturation.

Human neuromuscular organoids (NMOs) offer a powerful platform to overcome these limitations. We have previously demonstrated that self-organising human NMOs closely capture the developmental trajectory, tissue organisation and complexity of their in vivo counterpart, including the formation of morphologically and functionally competent NMJs that drive spontaneous contractions.^45^ Here, we hypothesized that introducing bioelectrical cues through controlled EPS training could further promote NMO maturation, acting as a physiological stimulus that is normally absent. To test this hypothesis, we developed a framework for low-frequency EPS training of NMOs and provided proof-of-concept that electrical stimulation can accelerate and enhance their functional maturation. Using acute stimulation assays and NMJ functionality as a readout, we first identified early NMO developmental stages as the optimum time window to initiate EPS training. While acute/short-term stimulation induced improvements in NMJ activity, these effects were transient, suggesting that prolonged stimulation is required to achieve stable and long-term maturation. We therefore established chronic EPS protocols, applying stable or dynamic pacing for approximately one month. Regardless of pacing parameters, chronic EPS elicited a significant boost in NMO maturation status, as evidenced by a dramatic increase in NMJ absolute number, size, and innervation levels, which translated in more frequent and stronger spontaneous contractions of EPS-NMOs. Strikingly, such enhanced activity persists for several days after training stops. Transcriptomic analysis revealed that chronic EPS, under stable or increasing pulse parameters, induced a significant upregulation of several genes involved in muscle contraction, myelination, and neuronal functions, consistent with accelerated neuromuscular maturation. Quantitative image analysis further substantiated these findings, revealing tissue-specific EPS-driven maturation effects. EPS-NMOs contained significantly higher number of glial cells, compared to non-paced control NMOs, with improved localisation and projection of motor neurons in the muscle region, which, in turn, exhibited significantly higher numbers of more elongated and better aligned myofibres. Importantly, mechanobiological characterisation assays confirmed that EPS-NMOs acquired significantly stiffer tissue properties and faster relaxation dynamics compared to non-paced control NMOs, consistent with more efficient ECC and force generation, and, thereby, stronger contractions. Overall, our findings establish chronic low-frequency EPS as a robust, non-invasive, and tunable approach to promote NMO maturation. Our study, not only showcases the benefits of integrating bioelectrical cues with organoid models, but it also highlights the importance of introducing such biophysical cues from early developmental stages, providing a flexible framework for guided, on-demand maturation of complex organoid models.

## Results

### Establishing an EPS training paradigm in NMOs

We have previously established a robust protocol for the generation of NMOs in 3D from neuromesodermal progenitors (NMPs), derived from various human pluripotent stem cell (hPSC) lines.^45^ A defining characteristic of 3D NMOs is the self-organisation of spinal cord neurons and skeletal muscle cells into functional neuromuscular junctions (NMJs). NMJs start forming around NMO developmental day 25-30, characteristic acetyl-choline receptor (AChR) clusters can be clearly detected by day 30, and by day 50 functional NMJs are formed, driving NMO spontaneous contraction.^45^ Based on this protocol, we derived NMPs from the human induced pluripotent stem cells (hiPSCs) KOLF2.1J^46^ (i.e., KOLF) and WTC-11, fluorescently tagged with mEGFP to target titin (TTN) protein (i.e., WTC^mTTNGFP^)^47^ and, subsequently, generated NMOs (**Figure 1a**).

**Figure 1:**
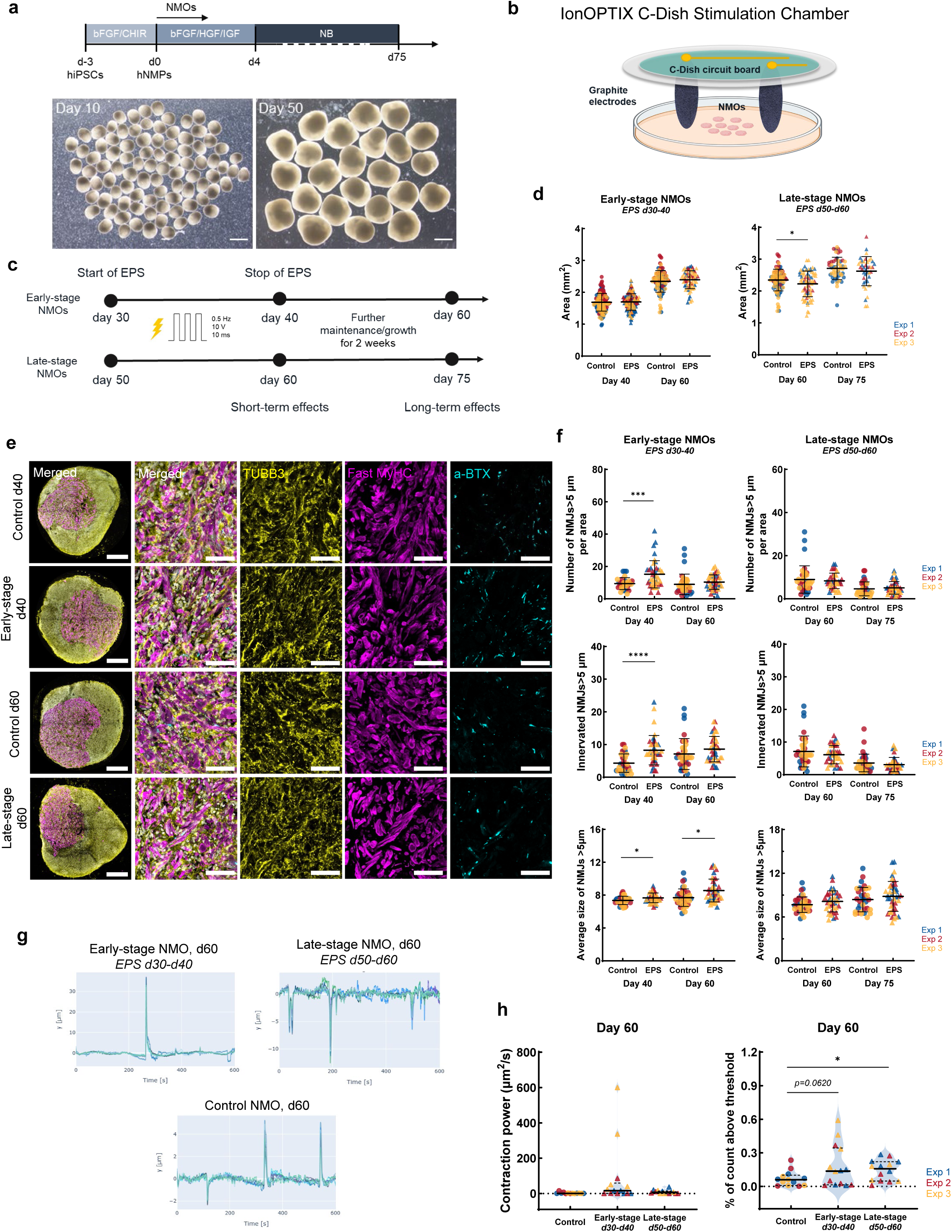
Defining the developmental window during which NMOs respond to EPS training. **a** Schematic representation illustrating the generation of NMOs from iPSC-derived NMPs. Representative brightfield images of day 10 and day 50 WTC^mTTNGFP^ NMOs. Scale bar 1 mm. **b** Schematic illustration of the EPS training experimental setup. For each pulsed stimulation training, NMOs are transferred to stimulation chambers comprising 6-well IonOptix C-Dishes and 6-well plates, prefilled with warm NB media. The schematic depicts one well of the stimulation chamber, with NMOs in the middle of the well, between a set of graphite electrodes, attached to the circuit board of the C-Dish. **c** Experimental design for identification of the NMO developmental stage at which EPS training can start. Day 30 or Day 50 WTC^mTTNGFP^ NMOs are exposed to AC stimulation with rectangular electrical bipolar pulses of 0.5 Hz, 10V, 10ms, for 30 min, every other day, for 10 days. At the end of this EPS training schedule, NMOs are collected for downstream analysis of the short-term EPS effects. Some NMOs are further maintained (day 60 and day 75) to evaluate potential long-term effects of EPS training. **d** Plots illustrating the size of Early-stage and Late-stage WTC^mTTNGFP^ NMOs, calculated using brightfield images and expressed as Area (mm^2^). The mean ± SD is shown for each experimental group (control in dots, EPS in triangle). Each datapoint represents one NMO. Data from *N=3* independent experiments are analysed by unpaired t-test with Welch correction (Early-stage NMOs: day 40 control *n=191,* EPS *n=141*, day 60 control *n=98,* EPS *n= 69*; Late stage NMOs: day 60 control *n=98,* EPS *n= 72; *P ≤ 0.05; **P ≤ 0.01; ***P ≤ 0.001; ****P ≤ 0.0001)*. **e** Confocal images of immunofluorescently labelled sections on WTC^mTTNGFP^ NMOs for neuronal (tubulin-β-3; TUBB3, in yellow), muscle (Fast Myosin Heavy Chain; MyHC, in magenta) and NMJ biomarkers (a-bungarotoxin; a-BTX in cyan), counterstained for DAPI (in grey). Representative whole NMO images are shown for each experimental group, at each timepoint of interest, along with 63x fields of view of the respective NMOs. Scale bar: 250 μm for whole NMOs, 50 μm for 63x micrographs. **f** Quantification of NMJ features based on 63x micrographs in e. Plots depict the absolute number per area (i.e., per 63x micrograph), innervation and average size of NMJs>5 μm on day 40 and day 60 for Early-stage NMOs and on day 60 and day 75 Late-stage NMOs. The mean ± SD of *N=3* independent experiments is shown for each experimental group. Each datapoint represents one 63x micrograph (control: day 40 *n= 29* from 11 NMOs, day 60 *n=37* from 13 NMOs, day 75 *n=35* from 13 NMOs; Early-stage NMOs: day 40 *n=39* from 13 NMOs, day 60 *n= 39* from 13 NMOs; Late-stage NMOs: day 60 *n=35* from 12 NMOs, day 75 *n= 45* micrographs from 15 NMOs;). Data are analysed by unpaired t-test with Welch correction (*\*P ≤ 0.05; **P ≤ 0.01; ***P ≤ 0.001; ****P ≤ 0.0001)*. **g** Representative Time Series plots, illustrating the spontaneous contractile activity of WTC^mTTNGFP^ non-paced control NMOs and EPS-NMOs on day 60, during a 10-minute live recording. **h** Quantitative analysis of the spontaneous contractile activity of WTC^mTTNGFP^ non-paced control NMOs and EPS-NMOs on day 60. *Contraction power* plot illustrating the strength of contractions in of each sample and *% of count above threshold* violin plot as an index of NMO contractile activity. Each datapoint represents one NMO (i.e., one 10-minute live recording). Data from *N=3* independent experiments are analysed by unpaired t-test with Welch correction (control: *n= 13* NMOs, Early-stage: *n=14* NMOs, Late-stage: *n=14* NMOs; *\*P ≤ 0.05; **P ≤ 0.01; ***P ≤ 0.001; ****P ≤ 0.0001)*.

We then used these NMOs to explore the potential of EPS training in enhancing the maturation status and functionality of organoids. As NMJs are established and maturing between days 30-50, we hypothesized that the timing of EPS would influence how NMOs respond to electrical activity. We therefore compared two developmental windows of NMOs: i) Early-stage NMOs, day 30, corresponding to the onset of NMJ formation and ii) Late-stage NMOs, day 50, corresponding to the stage when functional NMJs are already established. To deliver EPS, we used a multi-channel pacing system, compatible with standard culture formats (**Figure 1b**), previously used for chronic electrical stimulation of cardiomyocyte cultures^31,48–51^ and for modelling exercise in vitro.^38,41,42,52–54^ We screened a range of stimulation pulse parameters, including frequency (0.2 – 3Hz), pulse width (2-10 ms), and voltage (2-30 V) to identify conditions that elicited robust, synchronous, physiological contractions on day 65 NMOs. Rectangular biphasic AC electrical pulses of 0.5 Hz, 10 V and 10ms reliably induce a synchronous NMO contractile response, followed by full relaxation between pulses (**Supplementary Movie 1)**. This suggested that EPS-training could be used to stimulate NMOs and induce synchronous muscle contraction.

### Electrical pulse stimulation modulates NMJ morphology and function in a time-dependent manner

Using these optimized parameters, we subjected WTC^mTTNGFP^ NMOs to acute/short training of 30 mins of EPS (0.5 Hz, 10V, 10ms) every other day for 10 days (**Figure 1c**). Early-stage NMOs were trained from day 30 to day 40 and analysed on day 40 and on day 60, and Late-stage NMOs were trained from day 50 to day 60 and analysed on day 60 and on day 75 (**Supplementary movies 2-6**), in order to determine the short- and long-term effects of EPS on NMO morphology and function across developmental stages.

First, we compared the size of EPS-NMOs with non-paced control NMOs. EPS training did not interfere with the growth of Early-stage NMOs (EPS between d30-d40), which followed the same growth pattern as non-paced control NMOs (**Figure 1d** and **Supplementary Figure 1a,c**), exhibiting a normal increase in size on day 40 and on day 60. In contrast, EPS training on Late-stage NMOs (d50-60) induced a transient reduction in NMO size at day 60, but this effect resolved by day 75, when trained and control NMOs had comparable sizes. (Figure 1d and **Supplementary Figure 1b,c**). In addition, to examine whether EPS-training affected cell viability, we assessed apoptosis in NMOs, using cleaved Caspase-3 (cCas-3) immunofluorescence analysis (**Supplementary Figure 2a**). Quantification of cCas-3 expression in the neural and muscle compartments showed no difference in the expected baseline apoptotic levels between control and EPS-NMOs, indicating that EPS-training did not affect NMO survival (**Supplementary Figure 2b**).

Next, we examined the effects of electrical stimulation on NMJ development and morphology by analysing the NMJ number, size, and innervation in Early-stage and Late-stage EPS-NMOs compared to non-paced control NMOs (**Figure 1e**). NMJs were evaluated at two timepoints; on the day EPS-training stops (day 40, day 60) and ∼ two-three weeks post-EPS training (day 60, day 75), in order to assess both short- and long-term effects, respectively. Early-stage EPS produced a consistent increase in NMJ number and innervation at day 40, confirmed across three independent differentiations (**Figure 1f** and **Supplementary Figure 2c**). However, this effect persisted post-EPS only for NMJs>2μm, whereas larger NMJs>5μm returned to control levels by day 60. Moreover, NMJ morphological analysis revealed that NMJs>5μm were significantly larger in Early-stage EPS-NMOs compared to controls at both day 40 and day 60, indicating a lasting positive effect on NMJ maturation. In contrast, Late-stage EPS-training did not induce any significant changes in the size and innervation of NMJs compared to non-paced control NMOs at both timepoints (Figure 1f and Supplementary Figure 2c).

To further investigate the effects of EPS-training on NMJ status, we shifted our focus to the functional output of NMOs, analysing their spontaneous contractile activity on day 60; a developmental stage associated with more mature and functional NMJs that support skeletal muscle contraction in NMOs. Representative *Time Series* plots revealed enhanced contractility in both EPS-trained groups, which exhibited contractions of larger displacement (i.e., *y* axis; maximum distance from the organoid border, corresponding to 0 μm), compared to non-paced control NMOs (**Figure 1g and Supplementary Movies 7-9**). This effect was especially pronounced in Early-stage NMOs, despite EPS having ended ∼three-weeks earlier, suggesting long-lasting functional benefits. Quantitative analysis of contractile features confirmed these observations, revealing an EPS-induced increase in the *Contraction power* (i.e., the area under the curve of the squared time series over the duration of the recording, reflecting the overall magnitude of the signal), as well as a boost in the number of contractions above threshold in both experimental groups, with a significant increase in Late-stage EPS-NMOs (**Figure 1h**). The improved contractile output of Early-stage NMOs on day 60 correlated with their significantly larger NMJs, whereas the effect on Late-stage NMOs occurred without further NMJ size increase, suggesting that a different mechanism is at play. EPS training during early NMO developmental window (d30-d40) is time-matched to NMJ formation onset and precedes spontaneous contractile activity (∼d50). In contrast, Late-stage NMOs have already reached such milestones when EPS is applied (d50-d60) and can respond independently of NMJ function (Supplementary Movies 5,6). Indeed, we found that adding curare (10μM) to block NMJ activity does not affect the response of day 60 NMOs to EPS, which continue to respond to electrical pulses (1 Hz, 10V,10ms), exhibiting robust contractions despite NMJ blockage (**Supplementary Movie 10**).

Together, our results demonstrate that EPS training effectively modulates NMJ morphology and boosts functionality even after relatively short-term training (i.e., 10 days). However, the timing of training proved decisive for neuromuscular maturation, with early EPS-training coinciding with NMJ formation, yielding the most robust and enduring outcomes.

### Chronic electrical pulse stimulation enhances NMJ maturation status, independent of pulse parameters

Having identified early developmental stages as the optimal developmental window for effective NMO EPS-training, we then investigated how the duration of stimulation and the modulation of pulse parameters influence NMO maturation.

To this end, Early-stage NMOs (day 30) underwent one-month of chronic EPS training under i) stable parameters (i.e., Chronic EPS, *stable*: 0.5Hz, 10V, 10ms) or ii) progressively increasing parameters every 10 days (i.e., Chronic EPS, *increasing*: 0.5-1 Hz, 5-10V, 2-10ms), designed to follow the developmental trajectory of NMOs (**Figure 2a**). During this period, NMOs were subjected to EPS training for 30 mins, every other day, followed by analysis on day 60 (end of training) and on day 75 (two weeks post-training) to determine short- and long-term EPS effects, as before.

**Figure 2:**
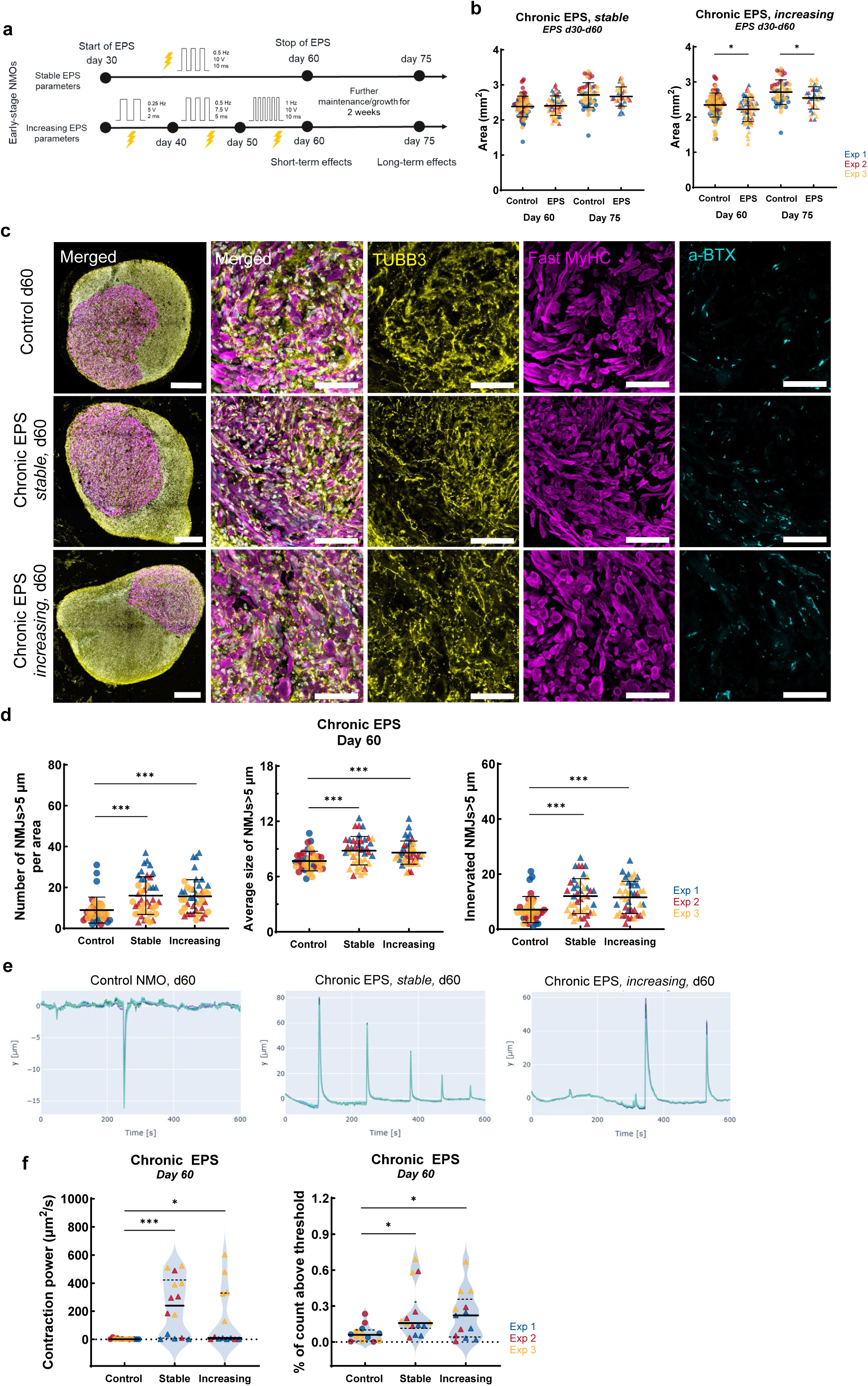
Modulation of NMJ function via chronic EPS training of Early-stage NMOs. **a** Schematic illustration of the modified experimental design for chronic EPS-training of NMOs. Early-stage (day 30) NMOs were subjected to EPS assays of either stable or increasing parameters, every other day, for ∼one month (day 60). NMOs were analysed on day 60 (short-term) and on day 75 (long-term) to evaluate the effects of chronic electrical stimulation on NMJ morphology and function. **b** Plots illustrating the size of WTC^mTTNGFP^ non-paced control NMOs and EPS-NMOs (expressed as Area (mm^2^)) on day 60 and day 75. Each datapoint represents one NMO. Data from *N=3* independent experiments are analysed by unpaired t-test with Welch correction (control: day 60 *n=86*, day 75 *n=51* NMOs; Chronic EPS, stable: day 60 *n=57,* day 75 *n=54* NMOs; Chronic EPS, increasing: day 60 *n=72,* day 75 *n=38* NMOs. *\*P ≤ 0.05; **P ≤ 0.01; ***P ≤ 0.001; ****P ≤ 0.0001)*. **c** Confocal images of immunofluorescently labelled sections of WTC^mTTNGFP^ NMOs for neuronal (tubulin-β-3;TUBB3, in yellow), muscle (Fast Myosin Heavy Chain; MyHC, in magenta) and NMJ biomarkers (a-bungarotoxin; a-BTX in cyan), counterstained for DAPI (in grey). Representative whole NMO images are shown for each experimental group, on day 60, along with 63x micrographs of the respective NMOs. Scale bar: 250 μm for whole NMOs, 50 μm for 63x micrographs. **d** Quantification of NMJs>5 μm features on day 60, based on 63x micrographs in c. Plots show the absolute number per area (i.e., per 63x micrograph), innervation and average size of NMJs>5 μm on day 60 for WTC^mTTNGFP^ non-paced control NMOs and EPS-NMOs, trained under stable or increasing pulse parameters. The mean ± SD of *N=3* independent experiments is shown for each experimental group. Each datapoint represents one 63x micrograph (control: *n= 37* from 13 NMOs, Chronic EPS, stable: *n=42* from 14 NMOs, Chronic EPS, increasing: *n=42* from 14 NMOs; Data are analysed by unpaired t-test with Welch correction (*\*P ≤ 0.05; **P ≤ 0.01; ***P ≤ 0.001; ****P ≤ 0.0001)*. **e** Representative Time Series plots, illustrating the spontaneous contractile activity of WTC^mTTNGFP^ non-paced control and EPS-NMOs chronically trained on day 60, during a 10-minute live recording. **f** Quantitative analysis of the spontaneous contractile activity on day 60 WTC^mTTNGFP^ NMOs. *Contraction power* plot illustrating the strength contractions in of each sample and *% of count above threshold* violin plot as an index of NMO contractile activity. Each datapoint represents one NMO (i.e., one 10-minute live recording). Data from *N=3* independent experiments are analysed by unpaired t-test with Welch correction (control: *n= 13* NMOs, Chronic EPS, stable: *n=14* NMOs, Chronic EPS, increasing: *n=13* NMOs; *\*P ≤ 0.05; **P ≤ 0.01; ***P ≤ 0.001; ****P ≤ 0.0001)*.

To assess if chronic EPS would elicit any changes in NMO growth trajectory, we quantified NMO size. NMOs exposed to stable stimulation parameters grew similarly to non-paced control NMOs. In contrast, those subjected to increasing parameters were significantly smaller at both day 60 (when EPS assay stops) and day 75 (two weeks post EPS; **Figure 2b** and **Supplementary Figure 3a,b**). Moreover, quantitative immunofluorescence analysis of cCas-3 expression in NMOs revealed that chronic EPS, regardless of stable or increasing pulse parameters, did not affect cell viability (**Supplementary Figure 3c,d**).

Next, we examined the effects of Chronic EPS on NMJ morphology and function using the same quantitative image analysis pipeline as before, across three independent differentiations. Strikingly, one month of EPS-training, applied under stable or under increasing parameters, induced a highly significant increase in the absolute number, size, and innervation of mature NMJs (>5μm) on day 60 compared to non-paced control NMOs (**Figure 2c,d**). However, this effect did not persist post-EPS on day 75 (**Supplementary Figure 4a,b)**. In contrast, intermediate NMJs (>2μm) showed a sustained increase in number, size, and innervation in both EPS-NMO groups, with a stronger effect on innervation in the stable parameter condition on day 60. However, this enhanced innervation persisted on day 75 only in the increasing parameter NMO group (**Supplementary Figure 4c)**.

Functional assays revealed that these morphological gains translated into improved contractile performance. Representative *Time Series* plots in **Figure 2e** clearly demonstrate a marked boost in NMO contractile activity upon one-month EPS training, with both EPS-NMO groups exhibiting considerably larger displacement, compared to non-paced control NMOs, as well as more contractions, especially in the Chronic EPS, *stable* NMO group (**Supplementary Movies 11-13**). Quantitative analysis further supports this, revealing significantly higher *Contraction power* and contraction *counts above threshold* in both EPS-NMO groups (**Figure 2f**). Importantly, such enhanced contractile function in EPS-NMOs was sustained two weeks post-training (**Supplementary Figure 5a,b and Supplementary Movies 14-16**), highlighting the positive and persistent effects of such physical conditioning.

Notably, one-month EPS training under stable parameters (i.e., Chronic EPS, *stable*: 0.5 Hz, 10V, 10 ms, for 30 mins, every other day) elicited the same benefit to the maturation status in an independent genetic background of KOLF NMOs. Both non-paced control NMOs and EPS-NMOs followed the same growth trajectory and comparable tissue segregation (**Supplementary Figure 6a,b**), but EPS-NMOs contained significantly higher numbers of intermediate NMJs (>2μm) and more mature NMJs (>5μm), of significantly larger size, and markedly increased levels of innervation (**Supplementary Figure 6c,d**). These findings also explain the enhanced spontaneous contractile activity observed in KOLF EPS-NMOs, exhibiting higher *Contraction power* and *counts above threshold* compared to non-paced control NMOs, however these differences were not significant (**Supplementary Figure 6e,f**).

Taken together, our findings demonstrate that chronic EPS enhances NMJ maturation and functional neuromuscular outputs, without compromising normal NMO growth, tissue organisation, or cell viability. Importantly, such outcomes were achieved with both stable and progressive pacing, underlining the effectiveness of EPS in driving neuromuscular maturation.

### Transcriptional analysis reveals chronic electrical pulse stimulation-induced maturation of NMOs

External electrical stimulation has been shown to induce significant transcriptomic responses in neural and muscle tissue, associated with neuronal and synaptic function and motor activity^55^ and with muscle structure, myogenesis and contraction,^56–58^ among others.

To explore EPS effects on NMO transcriptomic profile, we performed bulk RNA sequencing on EPS-NMOs and non-paced control NMOs at two timepoints: on day 40, corresponding to Early-stage, acute EPS-training, and on day 60, corresponding to chronic NMO pacing, under stable and under increasing pulse parameters. Differential gene expression analysis of day 40 revealed significant upregulation of only four genes and significant downregulation of another four genes, indicating that short-term, low frequency, low-voltage EPS-training (d30-d40) may modulate the morphological and functional features of NMJs without inducing broad transcriptional changes. In contrast, on day 60 we detected significant upregulation of 46 genes in EPS-NMOs subjected to stable pacing and of 52 genes in EPS-NMOs subjected to dynamic pacing (29 of those genes were common in both groups), while in the later a significant downregulation of 4 genes was also detected. Functional enrichment analysis on day 60 correlated the upregulated genes with several cellular processes, in both neural and muscle NMO tissues, including neuronal and dendrite function, axon guidance, myelination, muscle contraction, and myogenesis (**Supplementary Figure 7a**).

In both groups, this was driven primarily by a marked increase in the expression of *ARHGEF15, COL5A3, COL4A1, and ITGA1* over time, which is more prominent from day 40 to day 60 (**Supplementary Figure 7b**). *ARHGEF15* is known for its implication in physiological spatiotemporal development and maturation of synapses.^59^ Genes encoding for collagen type IV and V and associated receptors (*COL5A3, COL4A1, ITGA1*) play an essential role in Extracellular Matrix structural support and intercellular communication,^60,61^ including ECM-orchestrated NMJ organisation and long-term maintenance, neuronal connectivity, as well as in myogenesis and satellite cell activation.^62^ Moreover, we found that the highly significant and positive fold-change *of ITGA1* expression was also linked to improved muscle contraction in both EPS-NMO groups, validating our earlier findings on chronic EPS-boosted NMO functionality. Notably, neuronal and axon functions, along with muscle contraction and myotube formation, were also enriched by significant positive fold-changes in the expression of other genes (e.g., *ANXA1, MYH9, EGR1, and COL4A2)* in EPS-NMOs trained under stable parameters (Supplementary Figure 7a,b), further explaining the prominent effect on NMO spontaneous contractile activity we observed in this group.

Overall, these findings indicate that EPS training can effectively drive neuromuscular maturation, mainly through enrichment of functional pathways, while preserving NMO identity. Additionally, differential effects of stable versus dynamic pacing parameters on EPS-NMO transcriptomic profile highlight the robustness and flexibility of this approach in controlling and guiding several cell/tissue properties and functions according to desired outcomes by simply tailoring the electrical pulse parameters.

### Dynamic maturation of NMO neural tissue during chronic electrical pulse stimulation training

Having established a robust framework for chronic EPS training of NMOs, we then shifted our focus to tissue-specific effects. First, we assessed neural maturation using immunofluorescence analysis on day 40 (Early-stage NMOs) and day 60 (Chronic EPS, *stable* and *increasing* NMOs).

To examine the effects of EPS on the proliferation of neuronal progenitors, we co-labelled samples for SOX1 and Ki67 and quantified their expression at pre-determined timepoints (**Supplementary Figure 8a**). As expected, over time and as NMOs grow, proliferation of neural progenitors decreased. Chronic EPS training further reduced the Ki67^+^ cell population in the neural compartment on day 60, which was significant only in the Chronic EPS, *increasing* group, while the trajectory of SOX1^+^ cell population remained similar to non-paced control NMOs (**Supplementary Figure 8b**).

To evaluate astrocyte maturation, we analyzed Glial fibrillary acidic protein (GFAP) expression.^63^ As clearly seen in representative micrographs in **Figure 3a** and **Supplementary Figure 9a,** GFAP^+^ cells increased over time, both in non-paced control NMOs and in EPS-NMOs, as expected. Specifically, on day 40, GFAP^+^ astrocyte clusters localised in the central region of the NMOs, close to the neural-muscle tissue interface, while on day 60, extensive GFAP^+^ cell network could be observed in the neural NMO compartment. In fact, quantification of GFAP^+^ area confirmed a time-dependent expansion of astrocytic populations (d40 vs d60) and showed a significantly greater increase in the neural regions of day 60 EPS-NMOs, subjected to both stable and progressive pacing, compared to non-paced controls (**Figure 3b** and **Supplementary Figure 9b**).

**Figure 3:**
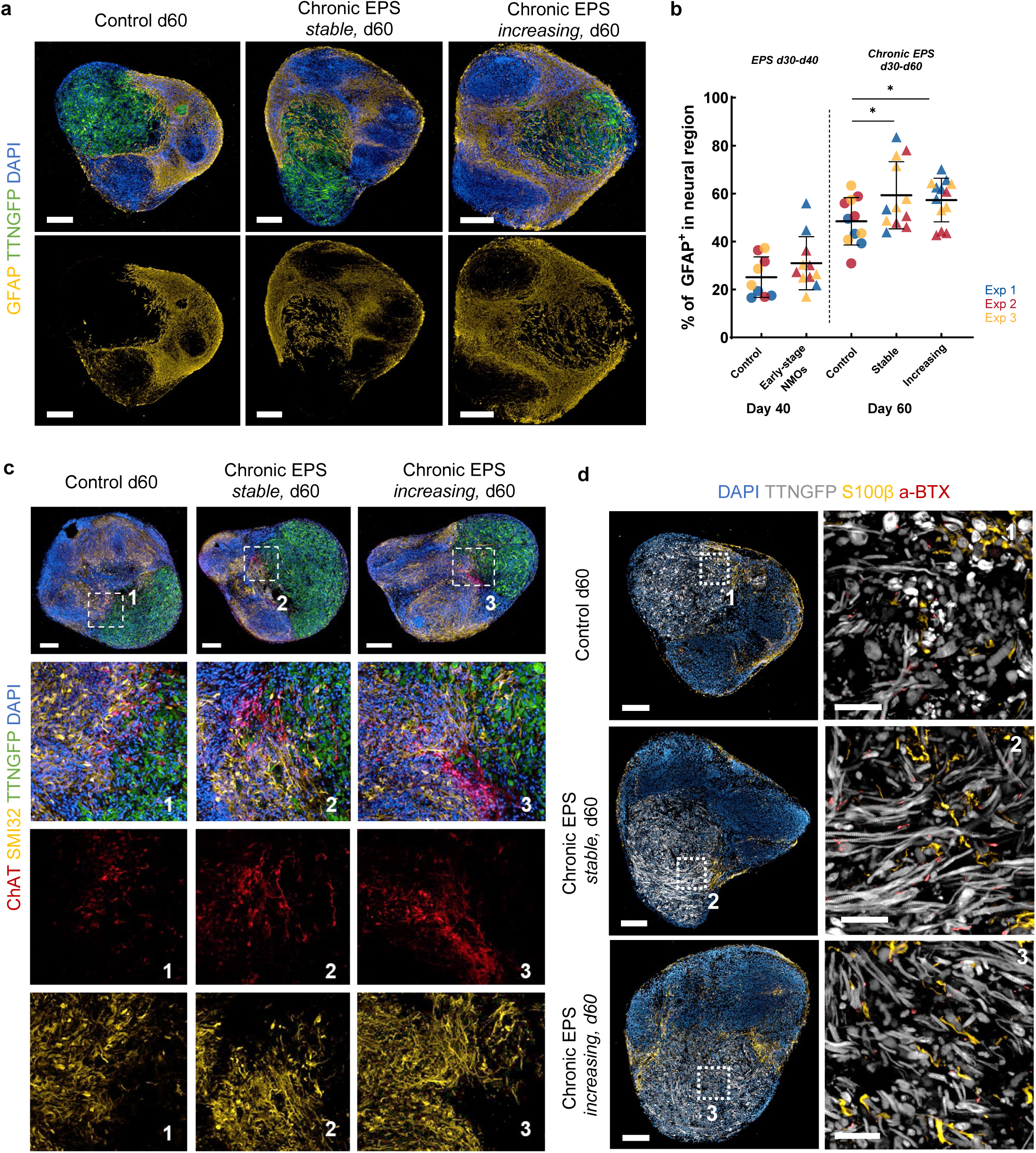
High-content, high-resolution imaging and evaluation of the EPS training effects on NMO neural tissue maturation. **a** Representative high-content, high-resolution images of whole NMO sections on day 60, illustrating the presence of glial fibrillary acidic protein positive cells (GFAP, in yellow) in WTC^mTTNGFP^ non-paced control and EPS-NMOs, trained chronically under stable or increasing pulse parameters. Scale bar 200 μm. **b** Quantification of the expression of GFAP^+^ glia cells in the neural area of WTC^mTTNGFP^ NMOs. The plot illustrates the GFAP^+^ cell expression over time in non-paced control NMOs and in EPS-NMOs on day 40 (Early-stage) and on day 60 (chronic EPS-NMOs, trained under stable or increasing pulse parameters). Each datapoint represents one NMO. Data from N=3 independent experiments are analysed by unpaired t-test with Welch correction (control: day 40 *n=9*, day 60 *n=11* NMOs; Early-stage: day 40 *n=11* NMOs; Chronic EPS, stable: day 60 *n=12* NMOs, increasing: day 60 *n=13* NMOs; *P ≤ 0.05; **P ≤ 0.01; ***P ≤ 0.001; ****P ≤ 0.0001). **c** Representative high-content, high-resolution images of whole WTC^mTTNGFP^ NMO sections (top panel) on day 60, depicting the presence of spinal cord motor neurons expressing the SMI32 neurofilament marker (in yellow) and/or the acetylcholine (ACh)-synthesizing enzyme choline acetyltransferase (ChAT; in red). Scale bar 200 μm. Bottom panels are higher magnification images of the respective annotated areas in whole NMO micrographs of the top panels, highlighting the organisation and localisation of SMI32+ with ChAT^+^ neurons, in regions where NMO neural and muscular segments interface. Scale bar 50 μm **d** Representative high-content, high-resolution images of whole NMO sections (left panel) on day 60, illustrating the presence of calcium-binding protein B positive Terminal Schwann cells (S100β, in yellow), in WTC^mTTNGFP^ non-paced control and in EPS-NMOs. Images in the right panel are close-ups of annotated areas in the muscle NMO region (TTNGFP, pseudo-coloured grey) of images in the left panel, revealing that S100β^+^ cells are present at very close proximity with NMJs (a-BTX, in dark red). Scale bar: left panel 200 μm, right panel 50 μm.

Next, by further leveraging high-content, high-resolution imaging, we examined the effects of EPS training on motor neuron maturation. We have previously demonstrated that NMOs contain spinal cord motor neurons expressing SMI32 neurofilament marker and Choline Acetyl Transferase (ChAT) enzyme, responsible for the biosynthesis of acetylcholine.^45^ EPS-NMOs exhibited stronger ChAT expression and clustering of ChAT^+^ motor neurons at the neural-muscle interface, as well as extensive expression of SMI32 by motor neurons that project axons in the skeletal muscle regions, consistent with an EPS-driven enhanced maturation status (**Figure 3c**).

Finally, we evaluated the expression of terminal Schwann cells as key components of healthy and functional NMJs. In line with our previous findings, we observed S100β^+^ terminal Schwann cell expression in close proximity with the aBTX^+^ AChR clusters at the NMJs. Careful observation of representative high-content, high-resolution immunofluorescence images revealed that S100β^+^ cells are highly localised at the neural-muscle region interface (**Figure 3d)**. This pattern appears more pronounced in both experimental groups, with more S100β^+^ neurite sheaths projected in the muscular regions, indicating axon and synaptic function enrichment, in line with our earlier observations.

### Dual role of chronic electrical pulse stimulation in skeletal muscle maturation

Electrical stimulation of skeletal muscle tissue in vivo and in vitro models is known to promote myoblast fusion and myogenic differentiation through satellite cell activation.^64,65^ To test whether EPS exerted similar effects in our study, we performed high-content, high-resolution quantitative image analysis of muscle progenitor and differentiation markers on day 40 and day 60 EPS-NMOs.

Co-labeling of NMOs for Paired Box 7 (PAX7) transcription factor and Ki67 proliferation marker revealed a highly proliferative PAX7^+^/Ki67^+^ cell population on day 60 EPS-NMOs under stable conditions (**Figure 4a,b** and **Supplementary Figure 10a,b**). Moreover, consistent with continuous NMO maturation, the expression of Myoblast Determination protein 1 (MyoD1), which regulates myoblast differentiation and maturation,^66^ decreased from day 40 to day 60. In line with prior literature,^67,68^ this trend was more pronounced in EPS-NMOs, which on day 40 already exhibited lower MyoD1 levels that continued to decrease and, by day 60, were significantly lower than in non-paced control NMOs (**Figure 4b** and **Supplementary Figure 11a**).

**Figure 4:**
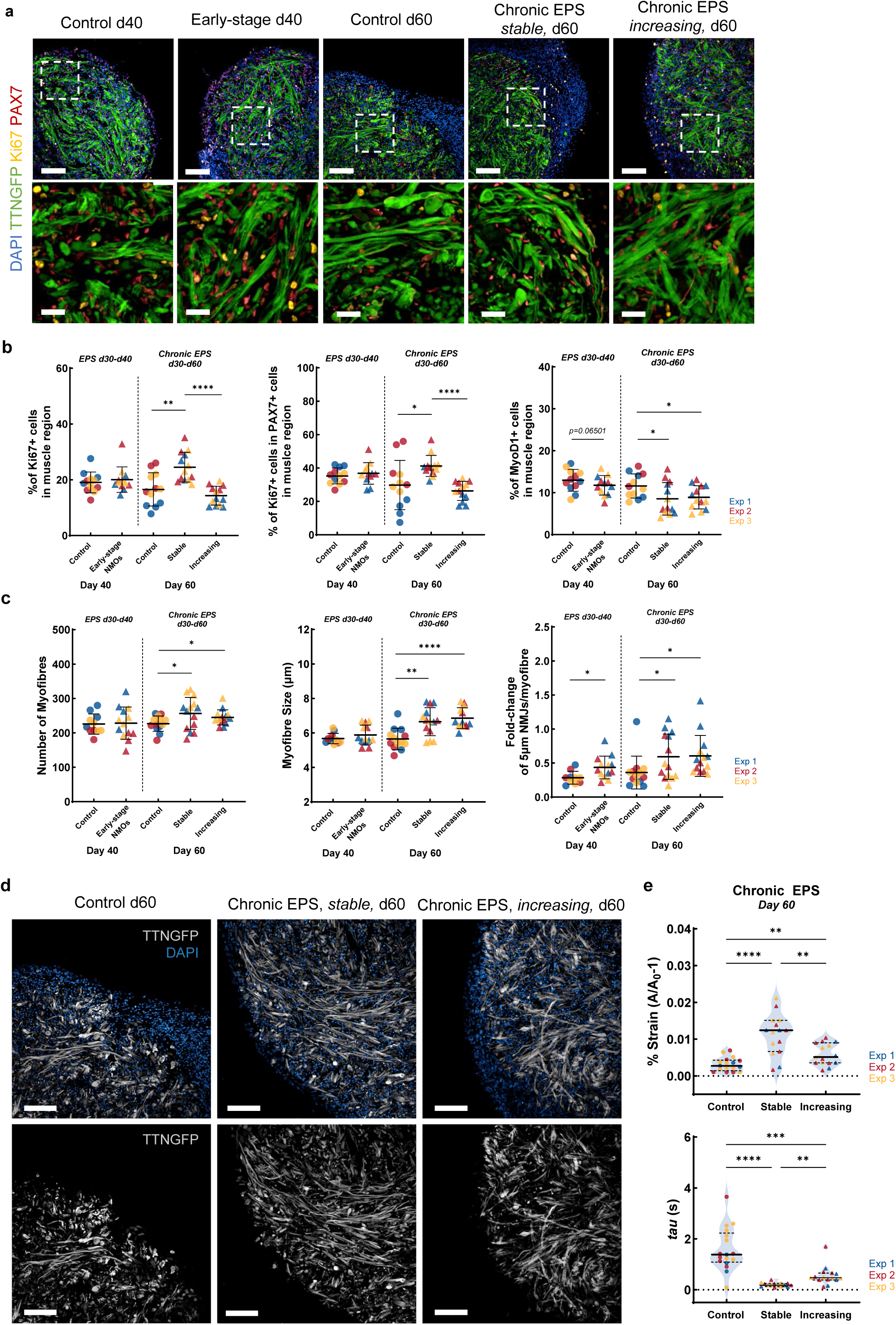
High-content imaging and quantitative analysis of the EPS training effects on NMO muscle tissue maturation. **a** Representative 20x high-content, high-resolution images (top panel) and respective magnified renderings (bottom panel). Magnified images in the bottom panel offer a close-up look in the NMO muscle fibre organisation and localisation with PAX7^+^ and/or Ki67^+^ cells in WTC^mTTNGFP^ non-paced control NMOs and EPS-NMOs. Scale bar: top panel 100 μm, bottom panel 25 μm. **b** Quantification of satellite-like (PAX7^+^), proliferating (Ki67^+^) cells, and myogenic progenitors (MyoD1^+^) in the NMO muscle region. Plots depict the changes in the expression of Ki67^+^ cells, Ki67^+^ in PAX7^+^ cells, and in MyoD1^+^ cells over time (d40-d60), between WTC^mTTNGFP^ non-paced control NMOs and EPS-trained NMOs (Early-stage on day 40 and Chronic EPS-NMOs, stable or increasing, on day 60). Each datapoint represents one NMO. Data from *N=3* independent experiments are analysed by unpaired t-test with Welch correction (control: day 40 *n=12*, day 60 *n=12* NMOs; Early-stage: day 40 *n=12* NMOs; Chronic EPS, stable: day 60 *n=13* NMOs, increasing: day 60 *n=13* NMOs; *P ≤ 0.05; **P ≤ 0.01; ***P ≤ 0.001; ****P ≤ 0.0001). **c** Quantification of muscle fibre features. Plots depict changes over time (d40-d60) in the absolute number of muscle fibres, in myofibre size, and in NMJ>5 μm number per myofibre in WTC^mTTNGFP^ non-paced control NMOs and EPS-trained NMOs (Early-stage on day 40 and Chronic EPS-NMOs, stable or increasing, on day 60). Each datapoint represents the mean number of fibres per three 63x micrographs and the mean size of muscle fibres per one 63x micrograph per NMO. The mean ratio of NMJ>5 μm/myofiber per three 63x micrographs was used for calculating the respective fold-change. Data from *N=3* independent experiments are analysed by unpaired t-test with Welch correction (control: day 40 *n=11*, day 60 *n=13* NMOs; Early-stage: day 40 *n=13* NMOs; Chronic EPS, stable: day 60 *n=14* NMOs, increasing: day 60 *n=14* NMOs; *P ≤ 0.05; **P ≤ 0.01; ***P ≤ 0.001; ****P ≤ 0.0001). **d** Representative high-content, high-resolution (20x) images of d60 WTC^mTTNGFP^ NMO muscle region sections. Inherent expression of Titin protein (TTNGFP; pseudo-coloured grey) demonstrates the organisation of the muscle sarcomere with typical striation, as well as muscle fibre fusion, and alignment. DAPI nuclear staining also highlights fibre peripheral nucleation, and the presence of multi-nucleated fibres. Scale bar 100 μm. **e** Biomechanical properties of WTC^mTTNGFP^ EPS-NMOs on day 60, compared to non-paced control NMOs. Plot in the top panel illustrates the maximum change in muscle region area (*A*/*A_0-1_*) due to tissue deformation during contraction, expressed as % of Strain, acting as an index of contraction strength. Plot in the bottom panel depicts relaxation dynamics, expressed as relaxation time constant (*tau*), acting as an index of tissue stiffness. These parameters are extracted from time-lapse data of NMO induced-contractions by electrical stimulation. Data from *N=3* independent experiments are analysed by unpaired t-test with Welch correction (control: *n=17* NMOs; Chronic EPS, stable: *n=15* NMOs, increasing: *n=15* NMOs; *P ≤ 0.05; **P ≤ 0.01; ***P ≤ 0.001; ****P ≤ 0.0001).

We further examined the effects of EPS on NMO muscle formation and maturation, based on immunofluorescence analysis of Fast Myosin Heavy Chain (Fast MyHC) expression. Quantification of absolute muscle fiber number and size (fiber diameter) revealed a significant increase in both features on day 60 EPS-NMOs, under either stable or increasing pulse parameters, but not after early, acute/short-term stimulation (d30-d40; **Figure 4c**). However, when we assessed the number of mature NMJs (>5μm), normalised for the number of muscle fibres – a key marker during neuromuscular development – we found a significant progressive increase between day 40 to day 60, compared to non-paced control NMOs (Figure 4c). The trend was similar in newly-formed NMJs (>2μm) and consistent across both EPS-NMO groups (**Supplementary Figure 11b**), validating our early observations about enhanced NMJ morphology and function upon electrical stimulation training. Additionally, we evaluated the NMO muscle fiber morphology qualitatively, leveraging our NMO **TTNGFP^+^**inherent signal. **Figure 4d** illustrates representative 20x field-of-view micrographs of NMOs on day 60, derived from high-content, high-resolution imaging. What is strikingly evident in these images is the dramatic increase in the length of TTNGFP+ muscle fibers in the chronic EPS conditions compared to non-paced controls, indicating enhanced myotube formation.

Finally, we characterised the effects of chronic EPS training on the biomechanical maturation of NMOs by quantitatively analysing the strain index and relaxation time of stimulation-induced contractions during time-lapse recordings (**Supplementary Figure 12**). Both non-paced control samples and EPS-NMOs exhibited reproducible response to electrical pulses (1Hz, 10V, 10ms), as evidenced by muscle region contractions, followed by exponential relaxation toward baseline (**Supplementary movies 17-19**). However, after one month of training, both groups of EPS-NMOs exhibited significantly higher peak strain amplitude (% strain: *max peak A/A0-1*) and significantly lower *tau* values (faster relaxation after stimulation-induced contraction), compared to non-paced control NMOs (**Figure 4e**). The increase in the strain index suggests improved force transmission and ECC that support stronger NMO muscle contractions, which can be attributed to more mature muscle fibers with enhanced morphology and innervation, as shown earlier. The marked decrease in *tau* values in EPS-NMOs, which is more significant in EPS-NMOs trained under stable parameters, reflects a more rapid relaxation following contraction, which is consistent with the development of stiffer tissues, with enhanced viscoelastic properties. In contrast, non-paced control NMOs retained higher *tau* values, as expected from more compliant and less mechanically mature tissues. Together, these findings suggest that EPS-NMOs undergo biomechanical maturation, acquiring properties of stiffer tissues, capable of stronger and faster contractions.

Collectively, the morphological and biomechanical analysis of NMO skeletal muscle tissue further substantiates our findings on EPS-guided maturation, emphasising again the importance of the prolonged duration of EPS training. Importantly, in our study, the nature of electrical pulse parameters emerged as a key factor, indicating that stable vs dynamic pulse parameters can activate distinct signalling pathways (e.g., PAX7^+^ cell proliferative capacity/status and *tau* values), thereby, highlighting the versatility of this method in modulating and controlling tissue properties according to desired outcomes, on demand and in a non-invasive and non-destructive way.

## Discussion

Organoids, as highly human-relevant models, are increasingly applied in biomedical research, holding great promise in shifting the paradigm of fundamental and translational studies, disease modelling, and drug discovery. However, recapitulating adult tissue maturation stages in organoids has been particularly challenging. Significant progress has been made recently to address this challenge, in synergy with bioengineering approaches. Biomechanical loading and/or electrical stimulation have emerged as key drivers for functional maturation of various bioengineered tissues.^30,31,69,70^

Here, we established a framework for chronic, low-frequency EPS as a training paradigm for guided maturation of complex human NMOs. This framework allowed us to treat EPS not as a single perturbation, but as a systemic training paradigm, in which electrical pacing during distinct developmental windows was used to test how activity influences the trajectory of NMJ formation and function. Time-matching the delivery of EPS training to the onset of synaptogenic programmes in NMOs was proven decisive. We found that subjecting NMOs to electrical cues from early developmental stages has a more prominent effect on the NMJ maturation, supporting strong and enduring muscle contractions, even upon acute/short-term EPS assays. In contrast, no significant effects were observed on the NMJ network of ‘older’ NMOs exposed to the same pacing cues, despite the improvement of spontaneous contractile activity. These findings suggest a dual action: in early-stage NMOs, EPS training promotes synaptogenesis likely by enhancing the clustering of AchR available at the onset of NMJ formation,^71,72^ while at later stages when NMOs have reached more advanced NMJ maturation stages, EPS training can no longer alter synaptic formation and maturation. Rather, it may act independently of NMJ function, as NMOs respond to EPS even upon NMJ blockage, mainly boosting the ECC capacity of NMO skeletal muscle, or act in conjunction with the already established NMJ network via activation of AchR clusters, thereby strengthening the existing motor neuron-muscle connections.

The next factor we identified as a key determinant for effective NMO maturation is the duration of EPS training. Chronic low-frequency EPS for ∼one month boosted NMO maturation status, significantly modulating tissue-specific cellular processes and functions, including functional neuromuscular output. Such enhanced functional capacity correlated with upregulation of transcriptomic programmes associated with more mature neuromuscular outputs in EPS-NMOs, without compromising the identity of the organoids. Notably, this was achieved both under stable and dynamic pacing, albeit via activation of different transcriptomic pathways, indicating that, by tailoring EPS parameters, it is possible to guide tissues toward desired outcomes.

Quantitative image analysis further resolved tissue-specific EPS effects. In the neural compartment, electrical stimulation led to a significant increase in glial cell population,^73^ and enhanced the localisation of motor neurons at the neural-muscle region interface, consistent with synaptic support and enhanced maturation. In muscle tissue the effects of EPS were multi-fold. EPS-NMOs exhibited significantly lower levels of MyoD1, indicating the presence of more mature muscle fibres,^67,68^ compared to control NMOs. Consistent with more mature tissue states, myofibres in EPS-NMOs were significantly larger, more elongated and better aligned, also containing higher numbers of mature NMJs (i.e., more NMJ>5μm per myofibre). Moreover, in line with literature, chronic EPS under stable pulse parameters upregulated myogenesis,^64,65^ leading to NMOs with significantly more myofibres and higher levels of proliferative PAX7^+^ cells, reflecting our transcriptomic enrichment for ECM and myogenic pathways. These findings suggest that the pacing programme could inform studies on skeletal muscle regeneration.

Finally, we also found that EPS-NMO muscle tissue undergoes biomechanical maturation, acquiring properties of increased stiffness, capable of stronger contractions and faster relaxation, in line with studies of EPS-training of skeletal muscle models.^38,42,54^ These effects were more pronounced in EPS-NMOs trained under stable parameters and could be driven by enhanced ECM deposition, as also suggested by higher expression levels of genes encoding for collagen and laminin receptors detected in this group. Importantly, this analysis complements and further validates our earlier findings on the capacity of EPS-NMOs for stronger spontaneous contractions (i.e., *Contraction power*), showcasing the power of our live imaging-based, non-destructive, and automated quantification pipeline for evaluating NMO contractile function.

This work is a proof of principle study conducted in two different genetic backgrounds of human iPSC lines. Although key phenotypes were reproduced in both WTC^mTTNGFP^ and KOLF2.1J backgrounds, broader genetic diversity and replication across independent lines/donors will be required to establish generalisation of the method. Additionally, the current EPS training protocol is labor-intensive and not yet scalable to very large numbers of NMOs, constraining throughput, necessary for screening applications. Moreover, we relied on bulk RNA-seq rather than single-cell RNA-seq, therefore cell-type-specific programmess remain unresolved. Finally, while EPS induced transcriptomic changes consistent with maturation, these changes do not appear to be the primary drivers of the enhanced functional outputs observed, a point that will require targeted perturbations for disentangling transcriptional from structural and biomechanical contributions.

Overall, here, we show proof-of-evidence of efficient EPS-guided maturation of complex self-organising multi-tissue organoids. Our findings highlight the powerful potential of electrical stimulation in modulating tissue properties in a non-invasive and non-destructive manner. We anticipate that the versatility of this approach will enable its broader application, facilitating morphological and functional maturation of further complex organoid systems as highly human-relevant tools of human physiology and disease. To this end, our focus is now shifted to integrating more advanced bioengineered approaches, which will enable on-demand, multi-modal monitoring and stimulation of organoids with high spatiotemporal resolution, and, thereby, mechanistic understanding of the development of human neuromuscular system. Our efforts are also concentrated on integrating technological advancements for making EPS training assays compatible with automated, high-throughput, and more user-friendly protocols and pipelines.

## Methods

### Generation of neuromuscular organoids

#### Human induced pluripotent stem cell lines

WTC-11 (GM25256) hiPSCs, fluorescently tagged with mEGFP to target titin (TTN) protein (WTC^mTTNGFP^, AICS-0048, obtained from Allen Cell Collection^47^) and KOLF2.1J hiPSCs, (HPSI0114i-kolf_2, JIPSC001000, obtained from the Jackson Laboratory^46^; **Supplementary Table 1**) were used for the generation of neuromesodermal progenitors and neuromuscular organoids, following the protocol described by Martins et al (2020).^45^ Both cell lines have been QC tested and the respective certificates of analyses were obtained from the providers.^46,47^ hiPSCs were passaged at least twice before used for experiments in this study (WTC^mTTNGFP^ passage 47+4 and KOLF2.1J P3^2^ +8). hiPSCs and NMOs were tested monthly for mycoplasma. Here, we directly used cryopreserved NMPs of the same bank/batch and performed at least three independent differentiations of NMPs into neuromuscular organoids for WTC^mTTNGFP^ and one differentiation for KOLF2.1J. A full list of reagents and consumables is available in Supplementary Information.

#### Cryopreservation of NMPs

At the end of differentiation, NMPs were washed once with 1X PBS and incubated with Accutase™(Sigma-Aldrich) or TrypLE™ (Gibco, Thermo Fisher Scientific) for 5 minutes at 37°C to obtain a single-cell suspension. To stop the enzymatic reaction, DMEM/F-12 (Gibco, Thermo Fisher Scientific) was added and the cell suspension was transferred to a 15 ml Falcon tube. NMPs were centrifuged at 2000 rpm for 4 min at room temperature, the supernatant was removed, and the pellet was resuspended in 1 mL N2B27. Cell counting was performed using Trypan blue (Gibco, Thermo Fisher Scientific) exclusion assay with Countess 3 automated cell counter (Thermo Fisher Scientific). Upon determining the volume of cryopreservation media required, NMPs were resuspended in Bambanker™ cryopreservation medium (Nippon Genetics) and aliquoted in cryovials. After overnight freezing at –80°C, cryovials were transferred to liquid nitrogen for long-term storage.

#### Generation of NMOs

On day 0, cryovials containing cryopreserved NMPs were briefly incubated in a 37°C water bath until just thawed. NMPs were immediately transferred to pre-warmed DMEM/F-12, followed by centrifugation at 2000 rpm for 4 minutes. The supernatant was removed, the cells were resuspended in N2B27 medium, and the cell concentration was determined upon counting, as described above. To generate NMOs, 9,000 NMP cells/well were plated in ultra-low binding 96-well plates (Thermo Fisher Scientific) in neurobasal medium (N2B27;100 μL/well; STEMCELL), supplemented with 50 μM Rho-associated protein kinase ROCK inhibitor (BioMol GmbH), 10ng/mL bFGF (produced in-house), 2ng/mL IGF1 (Peprotech) and 2ng/mL HGF (Peprotech). On day 2, 100 μL of fresh N2B27, supplemented with 2ng/mL IGF1 and 2ng/mL HGF, was added to each well of the 96-well U-bottom ultra-low attachment plates. On day 10, NMOs were transferred to 60mm dishes (Sigma-Aldrich; one dish per 96-well plate) with 6 mL of N2B27 and on day 20 each NMO dish was split in three 60mm dishes, again with 6mL N2B27 each. From day 4 no growth factors were added in N2B27 and media were replenished every other day, while from day 10, NMOs were maintained on an orbital shaker (75rpm). ^45^

### Electrical Pulse Training of NMOs

NMOs were included in EPS experiments upon quality checks at different developmental stages, including elongation and segregation of the NMO into mesodermal and neuronal compartments between developmental days 5-10. In addition, only NMOs that exhibited clear compartmentalisation at the predefined timepoints (day 30, 40, 60, 75) were selected for analysis.

For EPS pacing assays, each NMO experimental group was transferred to dedicated electrical stimulation chambers. Each chamber comprised a well of a 6-well plate, prefilled with 4.5 mL fresh N2B27 media, which was topped up to 6 mL by adding 1.5 mL of spent media from each NMO group dish, and a clean C-Dish graphite electrode circuit board (IonOptix Pace C-EM; IonOptix/CytoCypher B.V) placed between the well-plate and its lid. Plates were then transferred to the incubator and were connected to the IonOptix pulse generator dedicated channels, programmed to deliver pulses, according to the parameters of each experimental group (Figure 1c and 2a), for 30 mins. NMOs were then transferred back to their dedicated 60mm dishes, replenished with fresh N2B27 media.

The used C-Dish graphite electrode boards were then thoroughly cleaned, following the manufacturer’s protocol, and sterilised under UV for 30 mins before used again.

### Bulk RNA sequencing

#### Sample collection and RNA extraction

On day 30, 40 and 60, NMOs were collected for RNA sequencing by pooling and snap-freezing 3-5 NMOs per EPS group and controls. Total RNA per sample was extracted using the Direct-zol RNA Miniprep Plus Kit (ZYMO RESEARCH), following the manufacturer guidelines. Samples were stored at -80°C until ready for sequencing.

#### Library preparation and sequencing

Total RNA samples were quantified using a Qubit Fluorometer, and RNA integrity was checked on a TapeStation (Agilent). Double-indexed stranded mRNA-Seq libraries were prepared using the NEBNext Ultra II Directional RNA Library Prep with Beads Kit (NEB, # E7765L), starting from 400 ng of input material according to the manufacturer’s instructions. Libraries were equimolarly pooled based on Qubit concentration measurements and TapeStation size distributions. The loading concentration of the pool was determined using a qPCR assay (Roche, #7960573001). Libraries were then sequenced on the Illumina NovaSeq X Plus platform using PE100 sequencing mode using a 10B flow cell with the 200-cycle reagent kit, with a target of 30 million reads per library. Libraries were sequenced on an Illumina NovaSeq X Plus using a 10B flow cell with the 200-cycle reagent kit. The run was configured for paired-end, dual-indexed sequencing with the following cycle scheme: 101–10–10–117 (Read 1—Index 1—Index 2—Read 2). Instrument and reagent specifications for the NovaSeq X Series (including 10B/200-cycle kits) and the dual-index workflow are described by Illumina.

#### Read processing and quality control

Raw FASTQ files were assessed with FastQC (v0.12.1), and summary reports were inspected and collated with MultiQC (v1.25.1). Read-level quality metrics (per-base quality profiles, adapter content, duplication) indicated high-quality libraries with negligible adapter contamination; therefore, no read trimming was applied prior to alignment. FastQC documentation and modules were used as references for interpretation.

#### Alignment and quantification

Reads were aligned to the human reference genome GRCh38 (GENCODE v49/Ensembl 115 annotations) using HISAT2 (v2.2.1) with strand-specific settings (--rna-strandness FR) and otherwise default parameters. Aligned reads were sorted and indexed with samtools (v1.22.1). Gene-level counts were derived with featureCounts (Subread v1.5.3-0) in paired-end and stranded mode (-p -S 2) against the ensembl release-115 Homo sapiens Gene Transfer Format (GTF), producing a matrix of raw counts per gene per sample.

#### Differential expression analysis

Differential gene expression between EPS-stimulated and unstimulated conditions at days 40 & 60 was performed in R (v4.3.3) using DESeq2 (v1.46.0). Count data were imported without external normalization; DESeq2’s median-of-ratios method was used for size-factor normalization, followed by dispersion estimation and fitting of negative binomial generalized linear models. Unless otherwise specified, Wald tests were used for contrasts of interest, and P-values were adjusted for multiple testing using the Benjamini– Hochberg procedure. Genes with FDR < 0.05 and |log2FC| ≥ 1 were considered significantly differentially expressed.

#### Gene set enrichment analysis

Two complementary enrichment strategies were applied. (i) Over-representation analysis was conducted with enrichR (v3.4) using MSigDB Hallmark and GO Biological Process databases on significant up- and down-regulated genes (defined by the above-mentioned thresholds). (ii) Rank-based enrichment was performed with fgsea (v1.32.4) on preranked genes ordered by DESeq2 Wald statistic using 50000 permutations. Multiple-testing correction was applied to all enrichment results (Benjamini–Hochberg), and pathways with FDR < 0.05 were considered significant.

### Analysis of NMO Size

Before subjecting NMOs to EPS (day 30, day 50) and at pre-determined sample collection timepoints, 0.7x brightfield images of NMOs were acquired using a stereoscope (Olympus SZX16). For all analyses, only *.tiff images were used. Images were then processed using an automated pipeline to extract morphological features of the organoids. First, Cellpose, a widely-used framework for automated single-cell segmentation,^74^ was used to extract a segmentation mask of the organoids. Although Cellpose was originally developed to segment cellular structures, its use has been extended beyond single-cell segmentation to other round-like entities.^75,76^ Among the available models, the cyto 3 model was chosen here due to the more robust and accurate segmentation capabilities it offers, compared to earlier models, including image restoration support, denoising and blurring for enhanced border detection. In Cellpose, all images are by default resized so that all objects have approximately similar diameter (∼30 pixels). Given that our organoids are significantly larger, to enhance the accuracy of the model and guide it to efficiently detect objects at our desired scale, we defined an extra step for adjusting this default diameter, named *average diameter*. This value is determined by the user upon a quick evaluation of the NMOs with the largest and smallest diameter, based on each image scale bar. Subsequently, the image processing library for Python scikit-image^77^ was used to extract the area and the centroid coordinates from the segmented mask. The output of this analysis pipeline is a segmented image and an *.xlsx file (one per image), containing the individual label ID for each organoid, the organoid area in pixels, and the respective centroid coordinates. The user can manually evaluate the segmented image and make adjustments in the average diameter, if necessary, to improve segmentation accuracy or to exclude NMOs unsuitable for analysis. In this study, NMOs that did not exhibit distinct compartmentalisation in neural and muscle region were not included in downstream analysis. To quantify the NMO size, the area in pixels of each NMO is converted into physical units by the user, based on the scale bar of each image. This manual conversion, properly implemented in the excel file, allows use of the pipeline in other experimental setups or image formats, which may have different conversion factors, rendering the tool highly flexible and broadly applicable.

### Immunofluorescence analysis of NMOs

#### Sample collection, embedding and sectioning

At predefined timepoints for analysis, NMOs were fixed with 4% para-formaldehyde (PFA; VWR) for 1 hr on ice, followed by thorough washes with 1X PBS before overnight treatment in 30% sucrose. NMOs were then embedded using a warm 15% gelatin-10% sucrose (Sigma-Aldrich) solution, followed by solidification at 4°C and snap-freezing in isopentane before storing at -80°C. Organoids underwent cryosectioning in 16 μm thick slices (MicroM HM 560 Cryostat, Thermo Fisher), collected on Superfrost Plus microscope glass slides (epredia).

#### Immunofluorescence staining and imaging

Before staining, NMO sections were incubated in 1X PBS at 42°C for 3×30 mins to remove gelatin. Sections were then fixed with 4% PFA for 5 mins on ice, followed by permeabilization and blocking steps with 4% Bovine Serum Albumin (BSA; VWR) and 0.3% Triton X-100 (Sigma Aldrich) for 90 mins at room temperature, followed by overnight incubation with primary antibodies at 4°C (**Supplementary Table 2**). After thorough washing with 0.3% Triton X-100 (PBST, 3×30 mins), samples were incubated with species-specific secondary antibodies, conjugated with Alexa Fluor 488, 568, 647 (Supplementary Table 2), for 2 hr at room temperature, washed with PBST (3×30 mins), counter-stained with DAPI (Thermo Fisher Scientific), and mounted using Immu-Mount medium (epredia). Samples were then imaged using either a confocal microscope (Leica SP8) or a spinning disc confocal microscope (Opera Phenix Plus, Perkin Elmer).

#### Image analysis

Analysis of NMJs and myofibres was based on 63x confocal micrographs (LEICA SP8) of NMO sections co-labelled for a-BTX, Fast MyHC, and TUBB3. The absolute number of myofibres was quantified by manually counting the number of Fast MyHC^+^ myofibres in three 63x micrographs/per NMO, using the multi-point tool in FiJi ImageJ. Similarly, the size of Fast MyHC^+^ myofibre cross-sections was quantified in one 63x micrograph/per NMO as the minimum Feret’s diameter, using straight line ROI measurements in FiJi ImageJ. NMJ number, size, and innervation were analysed via a custom semi-automated image analysis pipeline of three 63x micrograph/per NMO, developed with ilastik (version 1.4.0.post1) and Python (version 3.9.21).The intensities of each a-BTX image were adjusted to a range of 10-84 in Fiji and saved as a new image. Ilastik models were trained for adjusted a-BTX and TUBB3 images. Both adjusted a-BTX and original TUBB3 images were segmented with ilastik, and the respective masks were exported as NumPy arrays (*.npy). These were then processed and analysed in Python, where a-BTX^+^ regions, attributed to NMJs, were used to calculate NMJ size and absolute number by resorting to the skimage library. NMJ number was calculated as absolute number per 63x micrograph. NMJ size was calculated as the length of the major axis of the ellipse that has the same normalized second central moments as the a-BTX^+^ region. TUBB3 masks were used to assess NMJ innervation, by expanding them by 2 pixels and overlaying with the corresponding NMJ mask. An NMJ was considered innervated if the a-BTX^+^ region overlaps with the expanded TUBB3 mask. Only NMJs>2µm were included in quantifications. The results of this analysis were automatically exported as output data *.xlsx files, along with confirmation images for validation and data analysis. The absolute number of NMJs per 63x micrograph was also normalised to the count of myofibres in the respective image. Fold-changes were obtained by dividing the NMJ number/myofibre calculated for each image at each time point (day 40, 60) by the mean NMJ number/myofibre of control images at the respective timepoint.

Spinning disc confocal images were used for the analysis of levels of apoptosis, proliferation, neural, and muscular tissue-specific progenitor and maturation marker expression in whole NMO sections, based on semi-automated custom image analysis pipelines developed in ilastik and Python. In all cases, muscle region was detected by Titin fibres, inherently expressing GFP (TTNGFP), unless otherwise stated. Prior to processing, masks of neural, muscle, and/or whole NMO were generated with FiJi, upon deconvoluting regions of interest (ROIs) based on tissue-specific biomarkers using the free-hand tool. In some images, GFP is pseudo-coloured white. Ilastik models were trained for several biomarkers of interest.

Apoptotic levels and glial cell population were quantified in whole NMO sections labelled for cCas-3 or GFAP, respectively, and counterstained with DAPI. In ilastik, cCas-3 and GFAP signals were segmented, and the respective masks were exported as NumPy arrays (*.npy). Upon processing in Python, apoptosis in neural and muscle regions was calculated as the percentage of cCas-3 area coverage in each region. Similarly, glial cell population was calculated as the percentage of GFAP^+^ area in neural and whole NMO regions.

Neural progenitors were quantified based on immunofluorescence signals of Ki67 and SOX1 in neural NMO compartment. In ilastik, images corresponding to each biomarker and to DAPI were segmented and the exported masks and *.npy arrays were processed in Python. SOX1 and Ki67 masks were filtered by the DAPI mask and the neural region mask. In the neural region, the percentage of SOX1+ cells was calculated as the area of SOX1^+^ signal divided by the area of DAPI^+^ signal. The percentage of Ki67^+^ cells was calculated similarly.

Muscle markers were quantified based on whole NMO sections labelled for MyoD1 and in sections labelled for PAX7 and Ki67, both counterstained with DAPI. Ilastik was again used to generate masks for each nuclear biomarker and for DAPI. In Python, MyoD1 and PAX7 masks were filtered by the DAPI mask and the muscle region mask. Expression of MyoD1^+^ and PAX7^+^ cell populations in muscle region were calculated as the area of the respective signal divided by the area of DAPI^+^ signal. Similarly, the percentage of Ki67^+^ cell population was calculated in the muscle region and in the whole NMO.

In all Python pipelines, results were exported as *.xlsx files, along with confirmation images, for validation and downstream data analysis.

### Functional characterization of NMOs

Spontaneous contraction of NMOs (day 60, 75) was recorded using a confocal microscope (Leica SP8), in incubation mode (37°C and 5% CO_2_). Two hours before the recordings, EPS-NMOs or non-paced control NMOs were collected and fresh N2B27 media was administered. One NMO per recording was placed in an empty well of the 12-well plate with minimum amount of N2B27 media (∼50 μL) in order to immobilise the sample and avoid drifting. With the 10x objective, a brightfield image of the sample was acquired and used as a map to determine location of the recording: either of the two outer edges of the NMO, where neural and muscle regions intercalate. Then, one 10-minute recording (6222 frames, 0.096 frames/s) of the chosen location is acquired using a 4.5x magnification, set such that roughly half of the frame is filled with the organoid. Live recordings from at least four NMOs from either control or EPS-trained organoids, from each independent differentiation and each independent genetic background were acquired.

To analyse the spontaneous contractile activity of NMOs from the above dataset, videos were processed in Python, upon developing a custom two-stage semi-automated analysis pipeline:

#### Signal extraction

During this stage, the movement of the organoid border over time is extracted as a *Time Series* signal. First, an intensity threshold, determined automatically using Otsu thresholding on the first video frame, is set to obtain a binary segmentation of the video. If necessary, this threshold is then adjusted manually by the user, before a binary segmentation of each video frame is generated. This is followed by a few morphological operations, to obtain the final binary mask, from which the organoid border is extracted. Next, the length of the organoid border is obtained for computing the minimum border length over the whole video, to ensure that the same border points are acquired throughout the recording.

Given that in each contraction recording, organoids have different orientation, a common point of reference for all organoid borders is required to measure their movement over time. To this end, each video frame is rotated by an angle (*θ*) either clockwise or counter-clockwise, such that the organoid border is always aligned to the *y*-axis. The rotated image is then cropped using the previously computed minimum border length. The resulting images include only the minimum border region, meaning the border region consistently visible across all frames. Finally, the border is split into 10-sub-regions, to ensure that localised twitches and subtle contractions will be included, and the average shift of the organoid border with respect to frame zero, (i.e., displacement of the organoid upon contraction) for each sub-region, is calculated. This results in extraction of 10 signals from each video, with length ***n*** equal to the number of frames in the video (6222 frames, 0.096 frames/s), visualised as *Time Series* plots.

#### Signal analysis

First, time series data undergo a pre-processing step, including polynomial interpolation of missing values (e.g., missing pixels in some border regions), conversion of time series units to physical quantities to match confocal settings (e.g., 1 pixel = 1.013 μm), noise reduction by smoothing, and data de-trending as a means to remove potential organoid drifting from the evaluation of the contractile activity. Contraction features are then extracted using specific features of interest from the Python package *tsfresh (Time Series FeatuRe extraction on basis of Scalable Hypothesis test)*^78^:

#### Contraction Power

To extract the absolute energy of our *Time series* signal, *y(t)*, which is the displacement (y-axis in *Time series* plots) of the NMO border over time, we used the Python package ts-fresh, in which the absolute energy of a signal is defined as the sum of the squared values of a signal, according to the following equation:

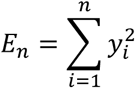

As this is often used a proxy of the overall magnitude of the signal across its duration, it was considered a suitable model for the analysis of our *Time series* signal, *y(t)*, thereby, defining *E_n_* as the area under the curve of the squared time series. However, this equation does not account for signal duration. To address this, we incorporated in the analysis the time resolution of each signal by dividing the absolute energy by the duration of the signal (i.e., *the number of bins, n, multiplied by the time resolution for a discrete signal, Δt*). We define this as *Contraction power*, computed according to the following equation:

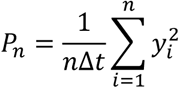

In this study, the time resolution of the signal is *Δt= 0.096 s*, corresponding to the sampling interval of the time series. Consequently, the units of the contraction power are μm^2^/s. Since the absolute energy represents the area under the squared signal, the contraction power represents the average value of the squared signal over time, acting as an indicator of the average intensity of the signal and, thereby, an indirect measure of the NMO contraction strength.

#### % of count above threshold

This feature describes the percentage of values in the time series (*y(t)*) that are higher than the given threshold (*th*) below which movements are considered noise. Empirically this threshold is set at 0.5 μm, but could be adjusted by the user if deemed necessary.

Finally, features are extracted for each contraction video as raw data in *.xlsx files for downstream analysis.

### Characterisation of NMO biomechanical properties

#### Video acquisition

NMOs were placed in the IonOptix stimulation chambers connected to the pulse generator (as described earlier). Contractions were induced upon EPS-stimulation of 1 Hz, 10V, 10ms. Time-lapse brightfield videos were recorded using a Leica DMi1 Stand microscope with FLEXACAM C1 and accompanying software (Enersight, Leica Microsystems), during EPS-induced contractions and subsequent relaxation time. All videos are of the same length (60 seconds, electrical pulses are delivered after the first 10 seconds) and were acquired under identical imaging conditions, before stored for further analysis.

#### Image processing and strain analysis

Analysis was performed for videos exhibiting the full dark region of the muscular region using Fiji. Organoid boundaries were identified using adaptive thresholding and morphological filtering the Particle Analyzer was used to extract the area. Python (PyCharm 2022.2.2) with OpenCV and NumPy was further used to plot A/A0 and extract peak strain as an index of contractile strength of the neuromuscular tissue. Relaxation dynamics (*tau*) were obtained by fitting the relaxation phase to an exponential decay model (**Supplementary Figure 12**). The relaxation time constant *tau* was interpreted as a measure of tissue stiffness and viscoelastic recovery. Lower *tau* values correspond to faster relaxation and mechanically stiffer organoids.

#### Data aggregation and Statistical Analysis

Strain parameters (peak strain, relaxation rate constants, residual strain) were summarized per organoid. Results across groups were aggregated over multiple experiments into a *.csv file for downstream analysis.

### Statistical analysis

Statistical significance was tested in GraphPad Prism 10. The number of biological and technical replicates, along with statistical tests used are indicated in figure legends. Significant differences are indicated as \**p ≤ 0.05; **p ≤ 0.01; ***p ≤ 0.001; ****p ≤ 0.0001*. Data are provided in a single Data Source File.

## Data availability

All analyses of sequencing data were executed on a SLURM-managed HPC cluster (Max Cluster) at MDC-Berlin using GNU parallel to parallelize QC, alignment, and counting across samples. Software versions and links to reference files (genome build and GTF) are provided in **Supplementary Table 3** and command-line arguments and analysis scripts are available on GitHub (https://github.com/Gouti-Lab/EPS-Project).

Image analysis was carried out using FiJi, iLastik and python scripts. Code is available on GitHub (https://github.com/Gouti-Lab/EPS-Project)

The code used for size analysis and spontaneous contraction characterisation in Python is available at https://github.com/HelmholtzAI-Consultants-Munich/NMOs-Contraction.

The programme for quantification of biomechanical properties is available at https://github.com/enricoklotzsch/organoid_stress_relax/

## Acknowledgements

The authors thank the genomics technology platform and pluripotent stem cell technology platform at the Max-Delbrück-Center for Molecular Medicine (MDC), Berlin, Germany, for technical support in this work. We also thank Andrea Grybowski and Noelle Findeisen for technical support.

## Funding

C-M.M wishes to acknowledge funding from the European Union Horizon 2023 research and innovation program under the Marie Skłodowska-Curie grant *e-NeuroMus* (Grant Agreement ID 101149182). M.G. wishes to acknowledge funding from the European Research Council (ERC) under the European Union Horizon 2020 research and innovation program (GPS-organoids; Grant Agreement No. 101002689). M.G. also receives funding from the European Molecular Biology Organization Young Investigator programme award and from the Einstein Stiftung Berlin (Einstein Center 3 R, EZ-2020-597-2). The work was supported by the Max Delbrück Center (MDC), which receives core funding from the Helmholtz-Association. E.K. acknowledges funding from the German Research Foundation (DFG; KL 3278/2-1 grant).

## Author contribution

M.G. and C.M-M. conceived the project and the experimental design. M.G. supervised the work, thoroughly reviewed and edited the manuscript, and provided critical feedback on data interpretation. C-M.M. designed and performed the experiments, collected all data and performed data analysis and interpretation, prepared the figures, and wrote the manuscript. I.A.M. established the pipelines for quantification of imaging data. I.A.E. analysed the bulk RNA sequencing data and prepared the respective figures. D.C, C.B, and I.M. established the automated analysis pipeline for organoid contraction properties and size quantification and M.P. supervised this work and provided feedback. I.L. generated and provided WTC^mTTNGFP^ NMPs. E.K. developed the pipeline and analysed the biomechanical properties of NMOs. All authors discussed the results and reviewed the manuscript.

## Ethics Declarations

## Competing interests

M.G. has filed patents for the method of human neuromuscular organoid generation. The remaining authors declare no competing interests.

## SUPPORTING INFORMATION

**Supplementary Figure 1:**
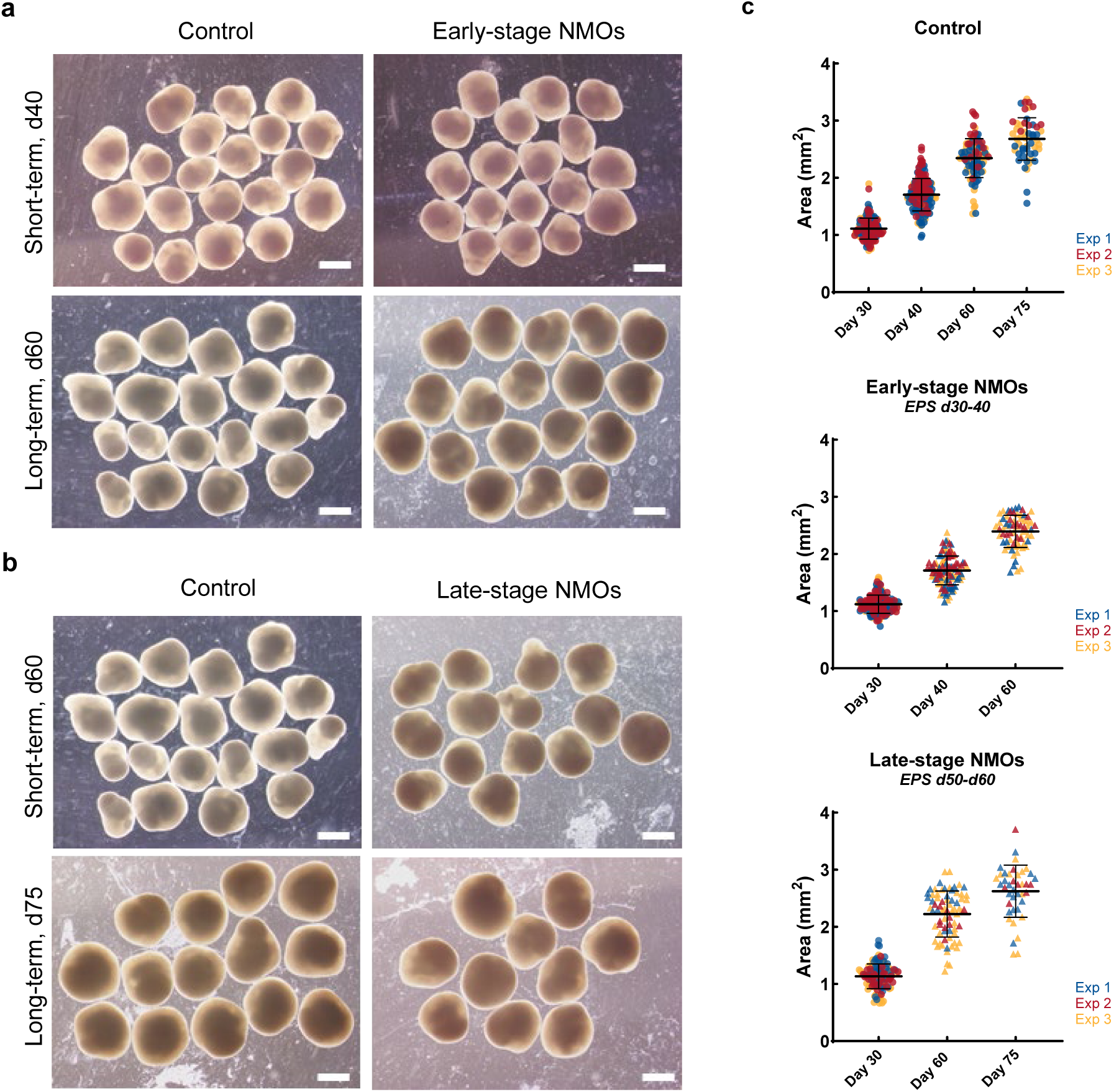
Supporting information for Figure 1d, related to WTC^mTTNGFP^ NMO Size. **a** Representative brightfield images of non-paced control NMOs and EPS-NMOs between day 30-40 (Early-stage NMOs), at different timepoints. Scale bar 1 mm. **b** Representative brightfield images of non-paced control NMOs and EPS-NMOs between day 50-60 (Late-stage NMOs), at different timepoints. Scale bar 1 mm. **c** Quantification of NMO size, calculated using brightfield images and expressed as Area (mm^2^). The mean ± SD is shown for each experimental group (control in dots, EPS-NMOs in triangle). Each datapoint represents one NMO. Data from *N=3* independent experiments are analysed by unpaired t-test with Welch correction (control: day 30 *n=159*, day 40 *n=170*, day 60 *n=98* NMOs; Early-stage: day 30: *n=153*, day 40: *n=109*, day 60 *n=69* NMOs; Late-stage: day 30 *n=166,* day 60 *n=72*, day 75 *n= 41* NMOs; *\*P ≤ 0.05; **P ≤ 0.01; ***P ≤ 0.001; ****P ≤ 0.0001)*.

**Supplementary Figure 2:**
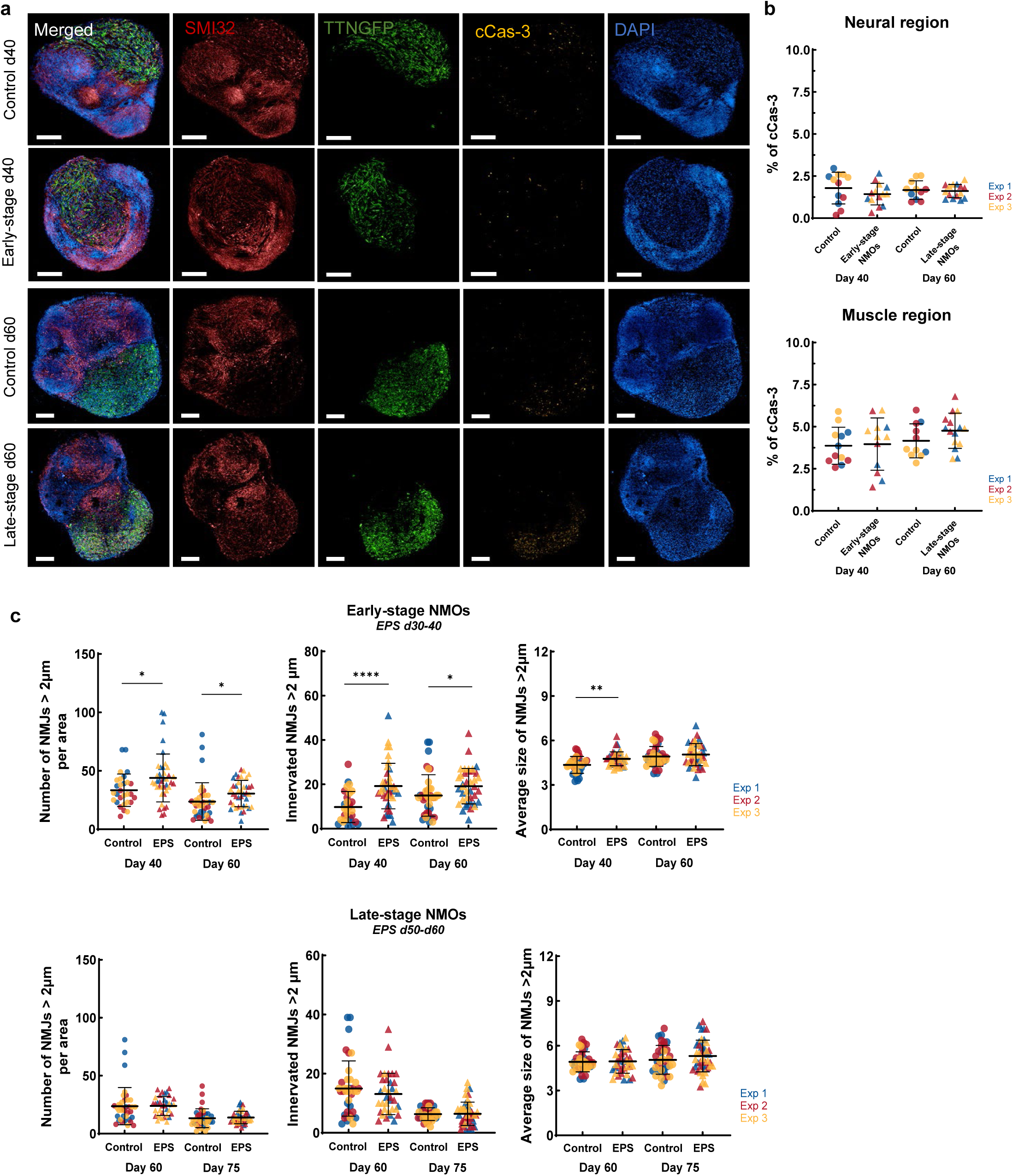
Evaluation of the EPS training effects on apoptosis levels in WTC^mTTNGFP^ NMOs. **a** Representative high-content, high-resolution images of whole-NMO sections of control, Early-stage and Late-stage EPS-NMOs, on day 40 and 60 (on the respective days EPS stops in both experimental groups), immunofluorescently labelled for the SMI32 neurofilament marker (in red), cleaved Caspase-3 protein (cCas-3; in yellow), and counterstained for DAPI. Titin protein (TTNGFP in green) is inherently expressed by NMOs. Scale bar 200 μm. **b** Quantification of Cas-3 apoptotic levels in neural and muscle NMO regions, based on immunofluorescence image data in a. Each datapoint represents one NMO. Data from *N=3* independent experiments are analysed by unpaired t-test with Welch correction (control: day 40 *n=12*, day 60 *n=11* NMOs; Early-stage: day 40: *n=12-14* NMOs; Late-stage: day 60 *n=15* NMOs; *\*P ≤ 0.05; **P ≤ 0.01; ***P ≤ 0.001; ****P ≤ 0.0001)*. **c** Additional quantification data on NMJ features, based on 63x confocal micrographs and focusing on NMJs>2μm. The mean ± SD of *N=3* independent experiments is shown for each experimental group. Each datapoint represents one 63x micrograph (control: day 40 *n= 29* from 1 NMOs, day 60 *n=37* from 13 NMOs, day 75 *n=37* from 13 NMOs; Early-stage NMOs, stable: day 40 *n=39* from 13 NMOs, day 60 *n= 39* from 13 NMOs; Late-stage NMOs: day 60 *n=35* from 12 NMOs, day 75 *n= 45* micrographs from 15 NMOs;). Data are analysed by unpaired t-test with Welch correction (*\*P ≤ 0.05; **P ≤ 0.01; ***P ≤ 0.001; ****P ≤ 0.0001)*.

**Supplementary Figure 3:**
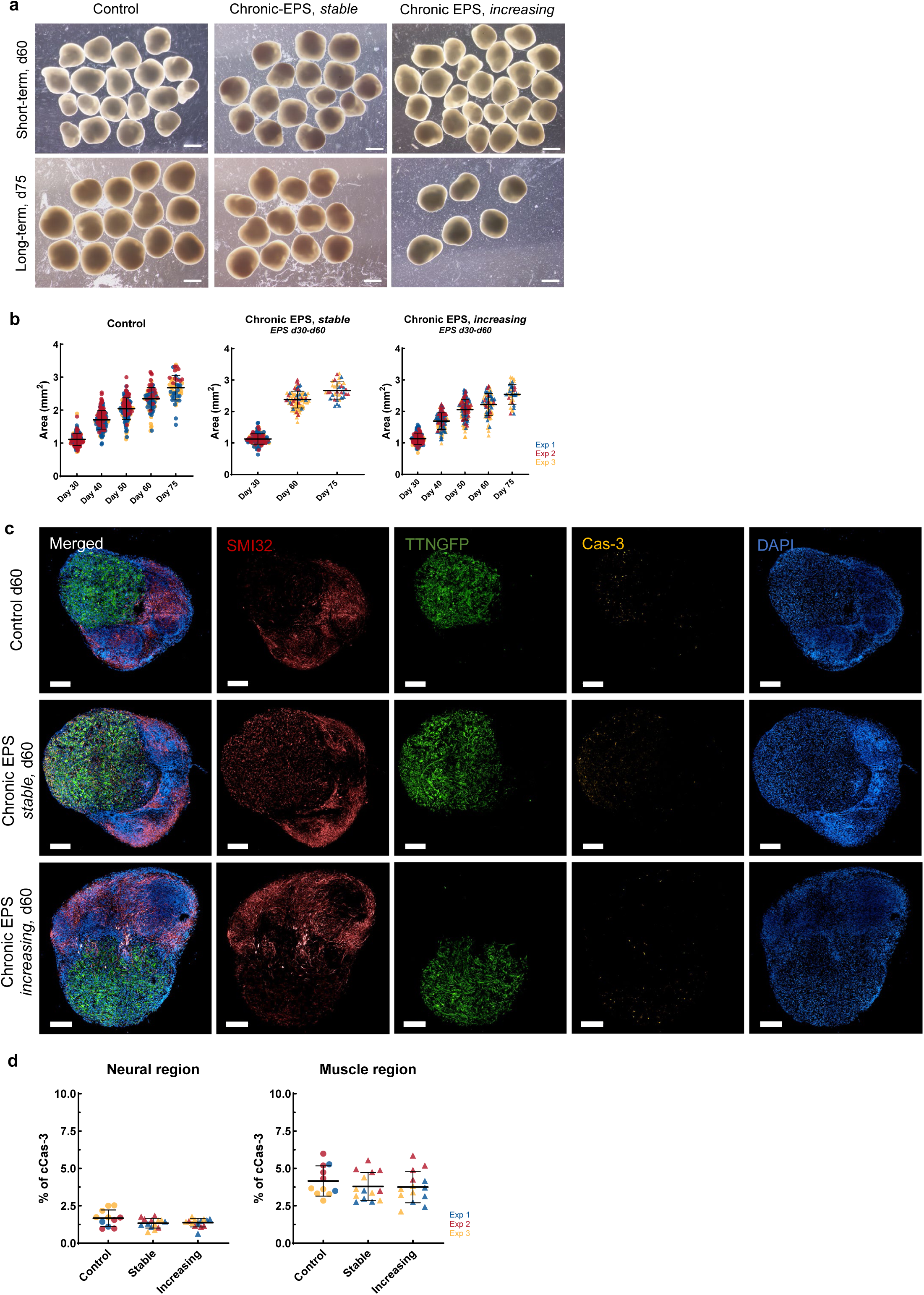
Supporting information for Figure 2b, related to EPS-effects on WTC^mTTNGFP^ NMO size and levels of apoptosis. **a** Representative brightfield images of non-paced control NMOs and EPS-NMOs, chronically trained between day 30-60, at different timepoints. Scale bar 1 mm. **b** Plots illustrating the growth of both non-paced control NMOs and EPS-NMOs over time, calculated using brightfield images and expressed as Area (mm^2^). The mean ± SD is shown for each experimental group. Each datapoint represents one NMO. Data from *N=3* independent experiments are analysed by unpaired t-test with Welch correction (control: day 30 *n=159*, day 40 *n=170*, day 60 *n=98* NMOs; Chronic EPS: stable day 30: *n=144*, day 60 *n=83,* day 75: *n=35* NMOs; increasing: day 30 *n=166,* day 60 *n=77*, day 75 *n= 38* NMOs; *\*P ≤ 0.05; **P ≤ 0.01; ***P ≤ 0.001; ****P ≤ 0.0001)*. **c** Representative high-content, high-resolution images of immunofluorescently labelled whole-NMO sections of non-paced control and EPS-NMO samples on day 60, when EPS training stops. Samples were labelled for the SMI32 neurofilament marker (in red), cleaved Caspase-3 protein (c-Cas-3; in yellow), and counterstained for DAPI (in blue). Titin protein (TTNGFP in green) is inherently expressed by NMOs. Scale bar 200 μm. **d** Quantification of day 60 cCas-3 apoptotic levels in neural and muscle regions of NMOs, based on immunofluorescence imaging data in c. Each datapoint represents one NMO. Data from *N=3* independent experiments are analysed by unpaired t-test with Welch correction (control: *n=11* NMOs; Chronic EPS: stable *n=14* NMOs, increasing *n=14* NMOs; *\*P ≤ 0.05; **P ≤ 0.01; ***P ≤ 0.001; ****P ≤ 0.0001)*.

**Supplementary Figure 4:**
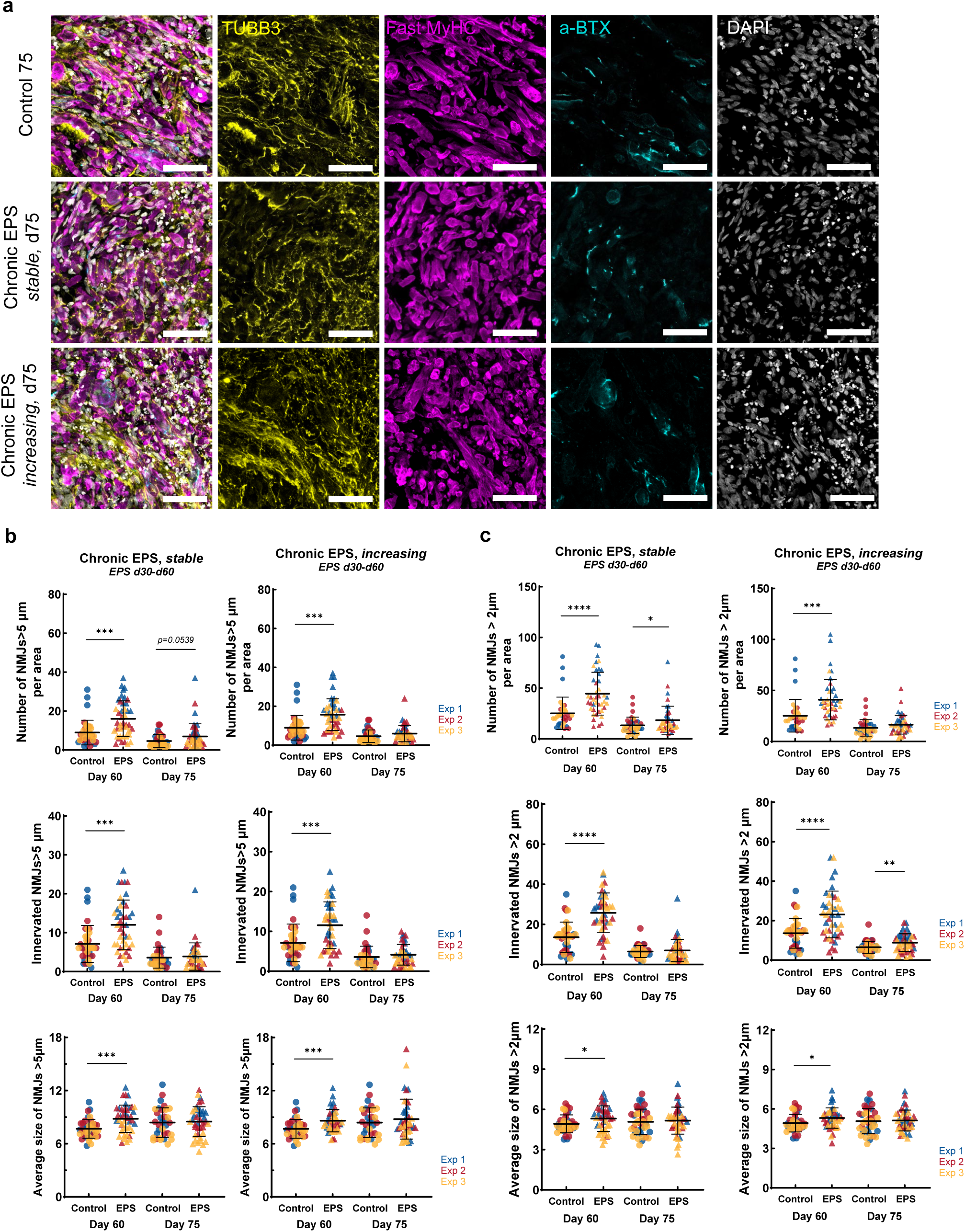
Supporting information for Figure 2c,d, related to EPS effects on NMJ features on day 75. **a** Representative confocal images (63x renderings) of immunofluorescently labelled sections of day 75 WTC^mTTNGFP^ non-paced control and EPS-NMOs, chronically trained until day 60. Samples were labelled for (tubulin-β-3;TUBB3, in yellow), muscle (Fast Myosin Heavy Chain; MyHC, in magenta) and NMJ biomarkers (a-bungarotoxin; a-BTX in cyan), counterstained for DAPI (in grey). Scale bar 50 μm. **b** Quantification of NMJs>5 μm features over time, based on 63x micrographs in b. Plots show the absolute number per area (i.e., per 63x micrograph), innervation and average size of NMJs>5 μm for non-paced control NMOs and EPS-NMOs, trained under stable or increasing pulse parameters, on day 60 (last day of EPS training) and on day 75 (two-weeks post-EPS training). The mean ± SD of *N=3* independent experiments is shown for each experimental group. Each datapoint represents one 63x micrograph (Day 60: control: *n= 37* from 13 NMOs, Chronic EPS: stable: *n=42* from 14 NMOs; increasing: *n=42* from 14 NMOs; Day 75: control *n=34* from 12 NMOs, Chronic EPS: stable: *n=38* from 14 NMOs, increasing: *n=42* from 15 NMOs; Data are analysed by unpaired t-test with Welch correction (*\*P ≤ 0.05; **P ≤ 0.01; ***P ≤ 0.001; ****P ≤ 0.0001)*. **c** Quantification of NMJs>2 μm features over time, based on 63x micrographs in b. Plots show the absolute number per area (i.e., per 63x micrograph), innervation and average size of NMJs>5 μm for non-paced control NMOs and EPS-NMOs, trained under stable or increasing pulse parameters, on day 60 (last day of EPS training) and on day 75 (two-weeks post-EPS training). The mean ± SD of *N=3* independent experiments is shown for each experimental group. Each datapoint represents one 63x micrograph (Day 60: control: *n= 37* from 13 NMOs, Chronic EPS: stable: *n=42* from 14 NMOs; increasing: *n=42* from 14 NMOs; Day 75: control *n=34* from 12 NMOs, Chronic EPS: stable: *n=38* from 14 NMOs, increasing: *n=42* from 15 NMOs; Data are analysed by unpaired t-test with Welch correction (*\*P ≤ 0.05; **P ≤ 0.01; ***P ≤ 0.001; ****P ≤ 0.0001)*.

**Supplementary Figure 5:**
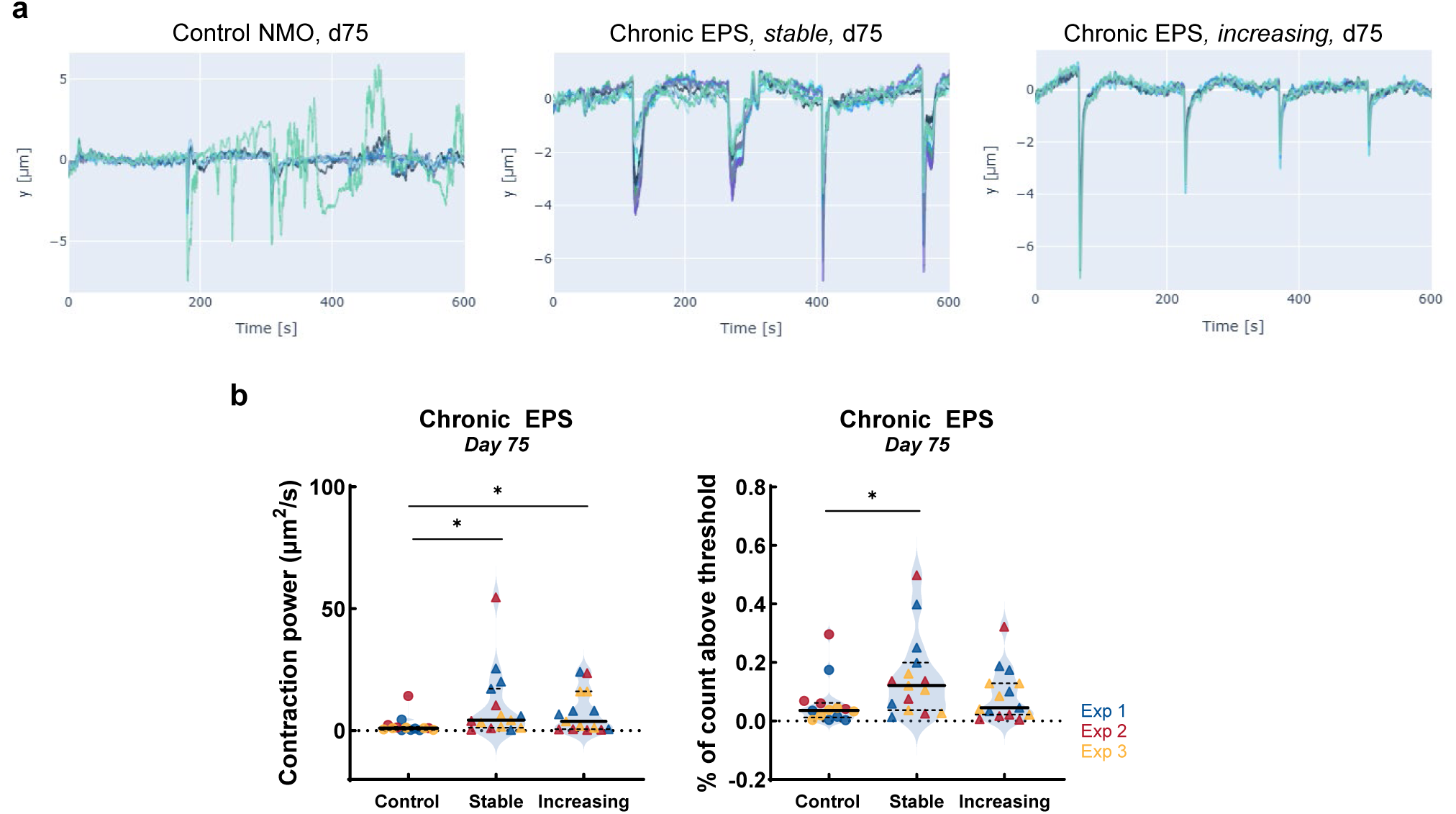
Evaluation of WTC^mTTNGFP^ NMO spontaneous contraction on day 75. **a** Representative Time Series plots, illustrating the spontaneous contractile activity of non-paced control and chronically EPS-trained NMOs on day 75, during a 10-minute live recording. **b** Quantitative analysis of the NMO spontaneous contractile activity on day 75, based on *Contraction power* (μm^2^/s) and % of count above threshold plots. Each datapoint represents the contractile activity of one NMO (i.e., one 10-minute live recording). Data from *N=3* independent experiments are analysed by unpaired t-test with Welch correction (control: *n= 14* NMOs, Chronic EPS, stable: *n=15* NMOs, increasing: *n=15* NMOs; *\*P ≤ 0.05; **P ≤ 0.01; ***P ≤ 0.001; ****P ≤ 0.0001)*.

**Supplementary Figure 6:**
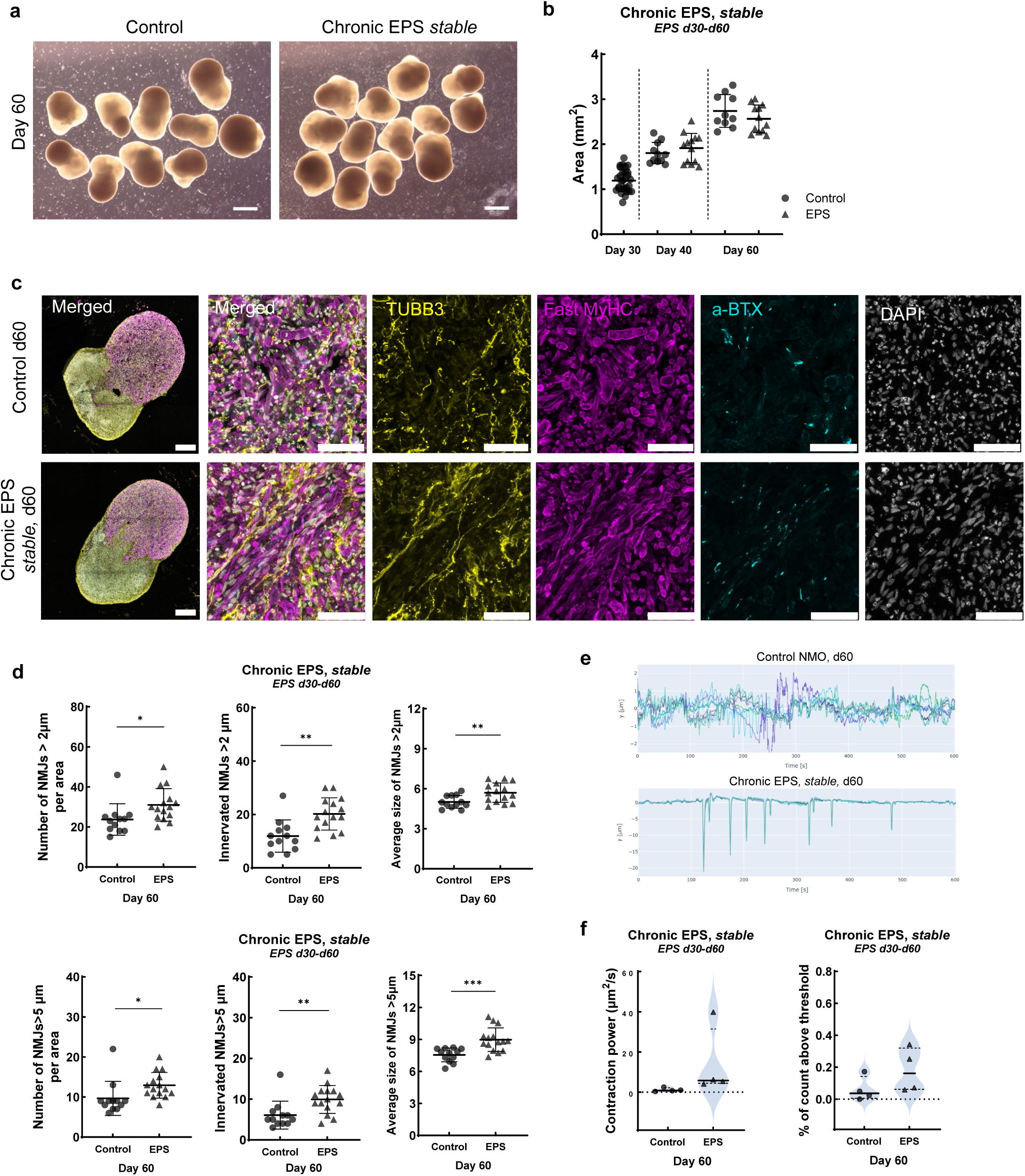
Evaluation of chronic EPS training under stable pulse parameters in KOLF NMOs. **a** Representative brightfield images of day 60 non-paced control KOLF NMOs and EPS-NMOs, chronically trained between day 30-60. Scale bar 1 mm. **b** Quantification of KOLF NMO size, calculated using brightfield images and expressed as Area (mm^2^). The mean ± SD is shown for each experimental group (control in dots, EPS-NMOs in triangle). Each datapoint represents one NMO. Data from *N=1* independent experiment are analysed by unpaired t-test with Welch correction (control: day 30 *n=29*, day 40 *n=11*, day 60 *n=10* NMOs; EPS-NMOs day 40: *n=13*, day 60 *n=12* NMOs; *\*P ≤ 0.05; **P ≤ 0.01; ***P ≤ 0.001; ****P ≤ 0.0001)*. **c** Confocal images of immunofluorescently labelled sections of day 60 KOLF non-paced control and EPS-NMOs, chronically trained until day 60. Samples were labelled for (tubulin-β-3;TUBB3, in yellow), muscle (Fast Myosin Heavy Chain; Fast MyHC, in magenta) and NMJ biomarkers (a-bungarotoxin; a-BTX in cyan), counterstained for DAPI (in grey). Representative whole NMO images are shown for each experimental group, along with 63x fields of view of the respective NMOs. Scale bar: 250 μm for whole NMOs, 50 μm for 63x micrographs. **d** Quantification of NMJ features on day 60, based on 63x micrographs in c. Plots show the absolute number per area (i.e., per 63x micrograph), innervation and average size of NMJs>2μm and NMJs>5 μm for KOLF non-paced control NMOs and EPS-NMOs, trained under stable parameters on day 60 (last day of EPS training). The mean ± SD of *N=1* independent experiment is shown for each experimental group. Each datapoint represents one 63x micrograph (control: *n= 12* from 4 NMOs, EPS-NMOs: *n=15* from 5 NMOs; Data are analysed by unpaired t-test with Welch correction (*\*P ≤ 0.05; **P ≤ 0.01; ***P ≤ 0.001; ****P ≤ 0.0001))*. **e** Representative Time Series plots, illustrating the spontaneous contractile activity of KOLF non-paced control and chronically EPS-trained NMOs on day 60, during a 10-minute live recording. **f** Quantitative analysis of the KOLF NMO spontaneous contractile activity on day 60, based on *Contraction power* (μm^2^/s) and % of count above threshold features. Each datapoint represents one NMO (i.e., one 10-minute live recording). Data from *N=1* independent experiment are analysed by unpaired t-test with Welch correction (control: *n= 4* NMOs, EPS-NMOs, stable: *n=4* NMOs; *\*P ≤ 0.05; **P ≤ 0.01; ***P ≤ 0.001; ****P ≤ 0.0001)*.

**Supplementary Figure 7:**
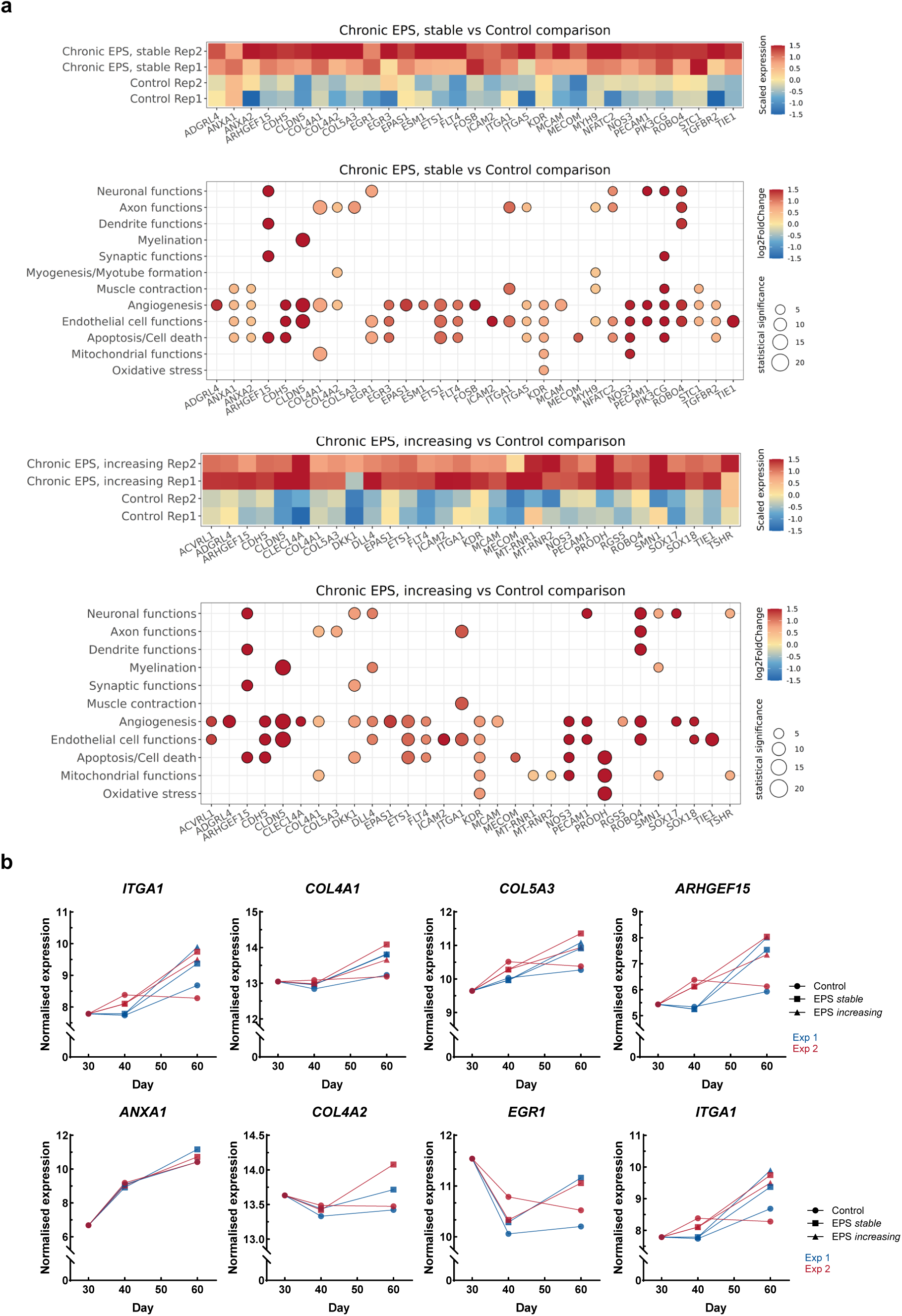
Analysis of WTC^mTTNGFP^ NMO transcriptomic profile. **a** Dot plots illustrating the differentially expressed genes between day 60 non-paced control NMOs and chronic EPS-NMOs (stable pulse parameters in top panel; increasing pulse parameters in bottom panel), mapped to the relevant biological functions and cellular processes they regulate. Dot colour indicates log₂ fold-change, and dot size represents the magnitude of statistical significance (*adjusted p-value <0.05*) **b** Normalised expression of DEGs (*adjusted p-value <0.05*) over time in non-paced control NMOs and EPS-NMOs. Top row plots depict the expression level trajectory of genes significantly enriched in both chronic EPS-NMOs, compared to non-paced NMOs. Bottom row panels illustrate the expression level trajectory of genes significantly enriched only in EPS-NMOs chronically trained under stable pulse parameters.

**Supplementary Figure 8:**
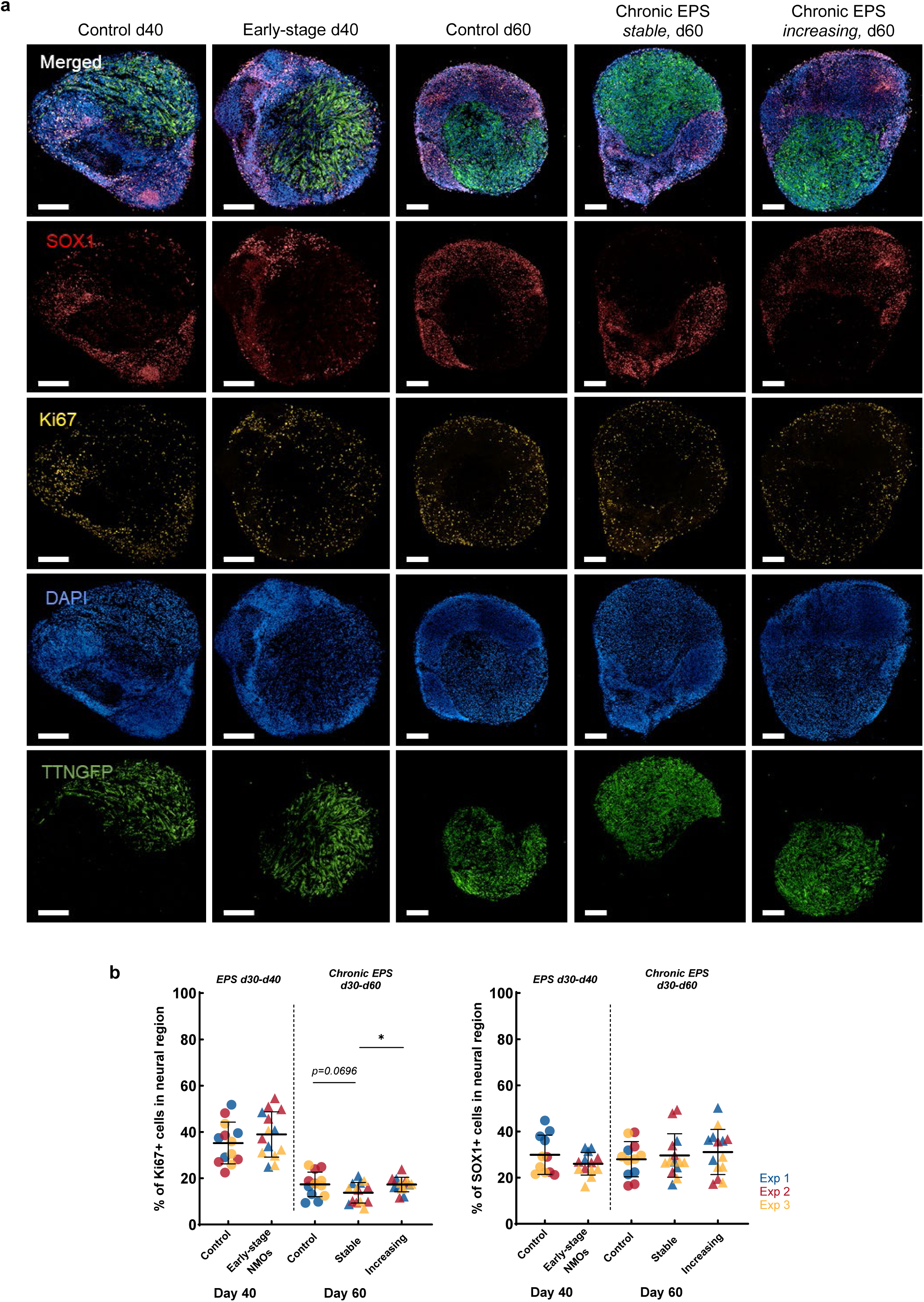
Image analysis of the neural progenitor and proliferation marker expression levels in WTC^mTTNGFP^ non-paced control NMOs and in EPS-NMOs. **a** High-content, high-resolution images of whole-NMO sections, immunofluorescently labelled for the neural progenitor marker SOX1 (in red) and proliferative marker Ki67 (in yellow), counterstained for DAPI (in blue). Titin protein (TTNGFP in green) is inherently expressed by NMOs. Images are representative of EPS-trained NMOs on day 40 (Early-stage) and day 60 (Chronic EPS-NMOs, under stable or increasing pulse parameters), and of non-paced control NMOs on the respective timepoints. Scale bar 200 μm. **b** Quantitative analysis of the expression of SOX1^+^ cells and Ki67^+^ cells in the neural region of EPS-trained NMOs on day 40 (Early-stage) and day 60 (Chronic EPS-NMOs, stable or increasing), and of non-paced control NMOs on the respective timepoints, based on images in a. Each datapoint represents one NMO. Data from *N=3* independent experiments are analysed by unpaired t-test with Welch correction (Day 40: control *n= 11-13* NMOs, Early-stage: *n=13-14* NMOs; Day 60: control *n=12-13* NMOs, Chronic EPS: stable *n=14* NMOs, increasing *n=14* NMOs; *\*P ≤ 0.05; **P ≤ 0.01; ***P ≤ 0.001; ****P ≤ 0.0001)*.

**Supplementary Figure 9:**
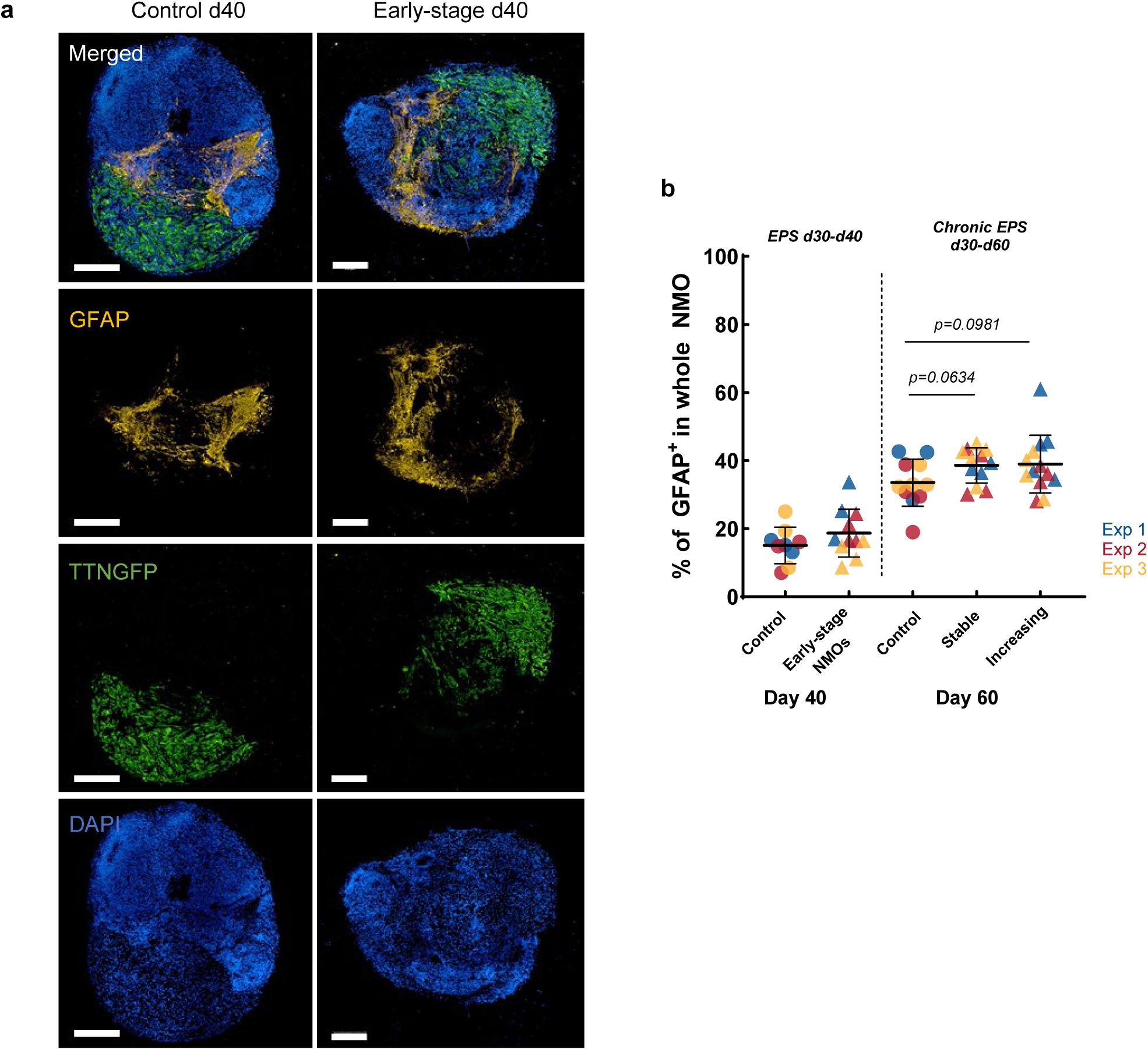
Supporting information for Figure 3, related to EPS effects on WTC^mTTNGFP^ NMO neural tissue cell composition and organisation. **a** Representative high-content, high-resolution images of whole NMO sections on day 40, illustrating the presence of glial fibrillary acidic protein positive cells (GFAP, in yellow) of non-paced control and Early-stage EPS-NMOs. Scale bar 200 μm. **b** Plot illustrating the expression of GFAP^+^ cells in the whole organoid of EPS-NMOs on day 40 (Early-stage) and day 60 (Chronic EPS, stable or increasing), and of non-paced control NMOs on the respective timepoints, based on images in Fig 4a. Each datapoint represents one NMO. Data from *N=3* independent experiments are analysed by unpaired t-test with Welch correction (control: day 40 *n=9*, day 60 *n=11* NMOs; Early-stage: day 40 *n=11* NMOs; Chronic EPS, stable: day 60 *n=12* NMOs, increasing: day 60 *n=13* NMOs; *P ≤ 0.05; **P ≤ 0.01; ***P ≤ 0.001; ****P ≤ 0.0001).

**Supplementary Figure 10:**
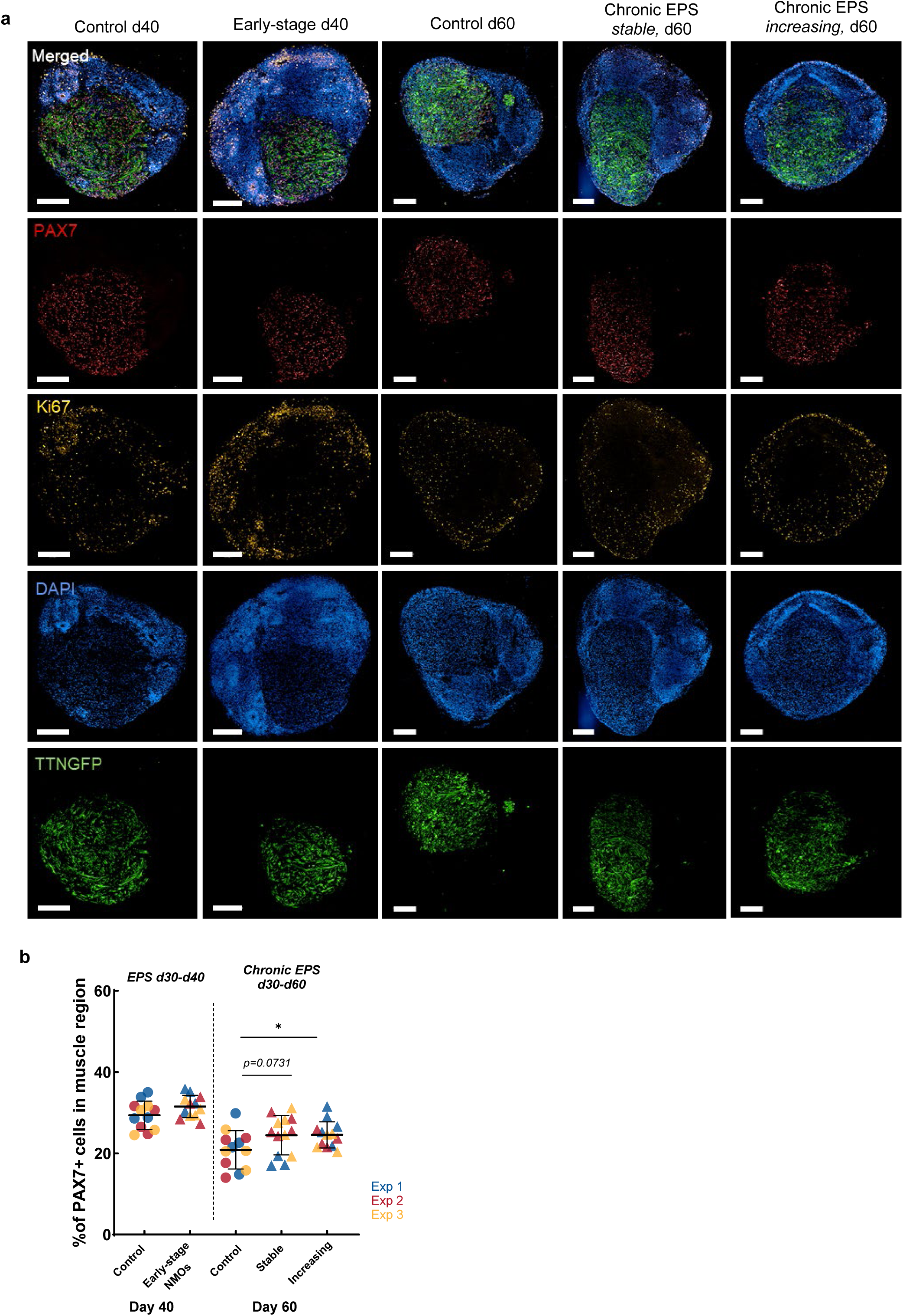
Image analysis of muscle progenitor and proliferation marker expression levels in WTC^mTTNGFP^ non-paced control and EPS-NMOs, related to Figure 4b, and quantification of muscle fibre features, related to Figure 4c. **a** High-content, high-resolution images of whole-NMO sections, immunofluorescently labelled for the satellite-like cell marker PAX7 (in red) and proliferative marker Ki67 (in yellow), counterstained for DAPI (in blue). Titin protein (TTNGFP in green) is inherently expressed by NMOs. Images are representative of EPS-trained NMOs on day 40 (Early-stage) and day 60 (Chronic EPS, stable or increasing), and of control NMOs on the respective timepoints. Scale bar 200 μm. **b** Plot illustrating the expression of PAX7^+^ cells in the muscle region of EPS-NMOs on day 40 (Early-stage) and day 60 (Chronic EPS, stable or increasing), and of non-paced control NMOs on the respective timepoints. Each datapoint represents one NMO. Data from *N=3* independent experiments are analysed by unpaired t-test with Welch correction (control: day 40 *n=12*, day 60 *n=12* NMOs; Early-stage: day 40 *n=12* NMOs; Chronic EPS, stable: day 60 *n=13* NMOs, increasing: day 60 *n=13* NMOs; *P ≤ 0.05; **P ≤ 0.01; ***P ≤ 0.001; ****P ≤ 0.0001).

**Supplementary Figure 11:**
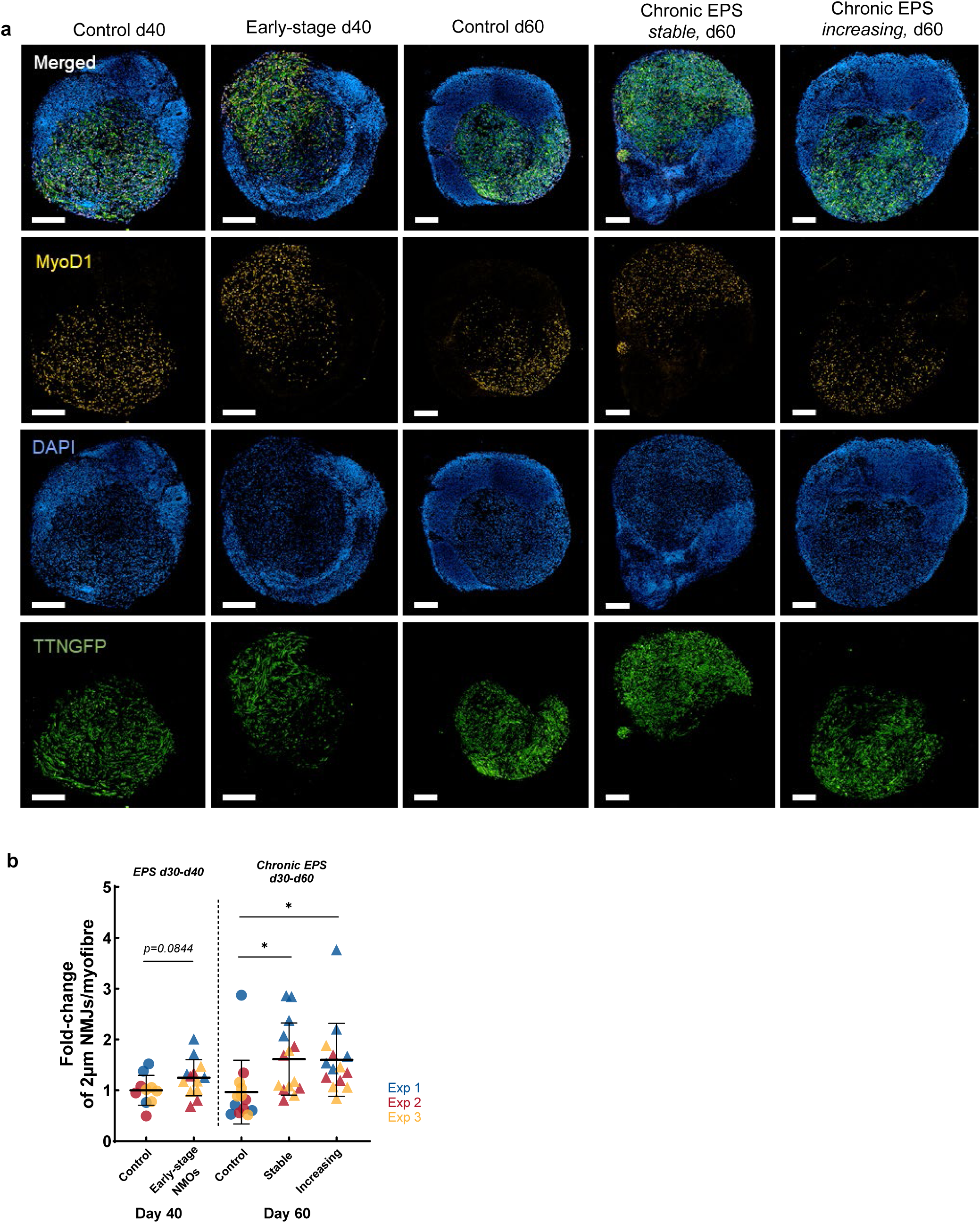
Supporting information for Figure 4b, related to evaluation of WTC^mTTNGFP^ NMO muscle progenitor levels. **a** High-content, high-resolution images of whole-NMO sections, immunofluorescently labelled for the myogenin progenitor marker MyoD1 (in yellow) and counterstained for DAPI (in blue). Titin protein (TTNGFP; in green) is inherently expressed by NMOs. Images are representative of EPS-NMOs on day 40 (Early-stage) and day 60 (Chronic EPS, stable or increasing), and of non-paced control NMOs on the respective timepoints. Scale bar 200 μm. **b** Analysis of the expression of NMJs>2 μm, normalised to the number of muscle fibres of sections of EPS-NMOs on day 40 (Early-stage) and day 60 (Chronic EPS, stable or increasing), and of non-paced control NMOs on the respective timepoints. Each datapoint represents the mean number of NMJs>2 μm/myofibre per two-three 63x confocal micrographs per NMO. Data from *N=3* independent experiments are analysed by unpaired t-test with Welch correction (control: day 40 *n= 29* from 10 NMOs, day 60 *n=37* from 13 NMOs; Early-stage: day 40 *n=39* from 13 NMOs; Chronic EPS, stable: day 60 *n=42* from 14 NMOs, increasing: day 60 *n=42* from 14 NMOs; *P ≤ 0.05; **P ≤ 0.01; ***P ≤ 0.001; ****P ≤ 0.0001).

**Supplementary Figure 12:**
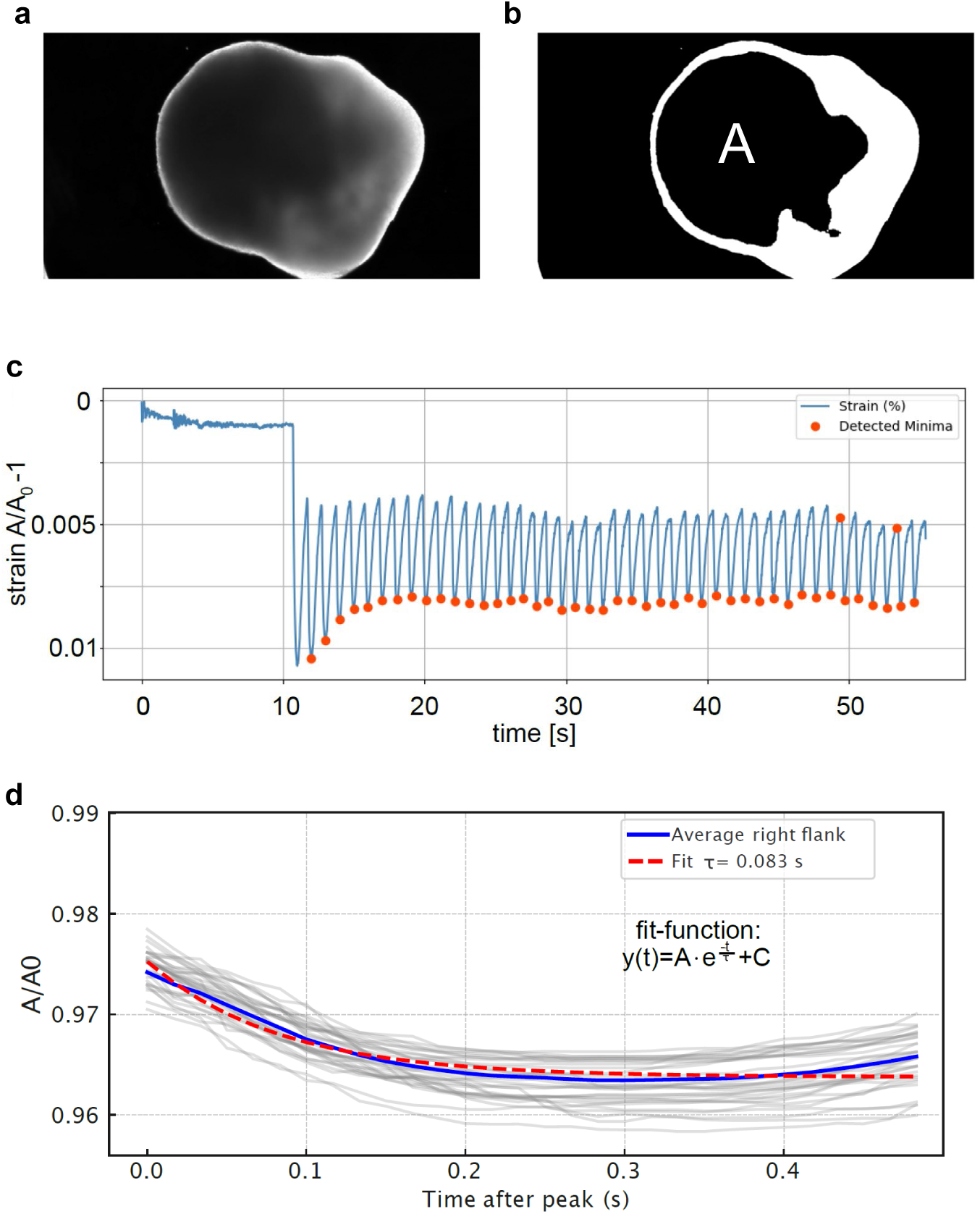
Supporting information for Figure 4e, related to evaluation of WTC^mTTNGFP^ NMO mechanical properties and respective analysis pipeline. **a** Brightfield image of an NMO, representative of a frame from one-minute live recordings of electrical stimulation-induced contractions. **b** Representative mask of the image in a, used for thresholding the muscle (dark region) of the NMO (*A*). **c** Plot illustrating the changes of muscle area (*A*) over time. Stimulation of the samples starts at 10 s and resumes until the end of the recording (60 s). Strain is calculated upon measuring the muscle area (*A*), normalised to the area in frame 0 (*A_0_-1*). *A* value of 0.01 represents a change in area of 1%. The maximum peak strain is calculated from the maximum and minimum of the first peak. **d** The plot illustrates the final step of the analysis pipeline for calculating the relaxation time constant *tau*. Individual curves from c are overlayed and fitted with an exponential decay model (shown in the plot), allowing for extraction of *tau*.

## Supplementary movies

Supplementary Movie 1: Representative video of a day 65 NMO during electrical pulse stimulation training (10V, 10 ms, 0.5 Hz)

Supplementary Movie 2: Representative video of a day 30 NMO response to electrical pulses (10V, 10 ms, 1 Hz)

Supplementary Movie 3: Representative video of a day 40 non-paced control NMO response to electrical pulses (10V, 10 ms, 1 Hz)

Supplementary Movie 4: Representative video of a day 40 Early-stage EPS-NMO response to electrical pulses (10V, 10 ms, 1 Hz)

Supplementary Movie 5: Representative video of a day 50 non-paced control NMO response to electrical pulses (10V, 10 ms, 1 Hz)

Supplementary Movie 6: Representative video of a day 60 Late-stage EPS-NMO response to electrical pulses (10V, 10 ms, 1 Hz)

Supplementary Movie 7: Representative video of a day 60 non-paced control NMO spontaneous contraction

Supplementary Movie 8: Representative video of a day 60 Early-stage EPS-NMO spontaneous contraction

Supplementary Movie 9: Representative video of a day 60 Late-stage EPS-NMO spontaneous contraction

Supplementary Movie 10: Representative video of a day 60 non-paced control NMO during electrical pulse stimulation and Curare (10 nM) exposure

Supplementary Movie 11: Representative video of a day 60 non-paced control NMO spontaneous contraction

Supplementary Movie 12: Representative video of a day 60 chronic EPS-NMO (stable) spontaneous contraction

Supplementary Movie 13: Representative video of a day 60 chronic EPS-NMO (increasing) spontaneous contraction

Supplementary Movie 14: Representative video of a day 75 non-paced control NMO spontaneous contraction

Supplementary Movie 15: Representative video of a day 75 chronic EPS-NMO (stable) spontaneous contraction

Supplementary Movie 16: Representative video of a day 75 chronic EPS-NMO (increasing) spontaneous contraction

Supplementary Movie 17: Representative video of a day 60 non-paced control NMO during electrical stimulation for biomechanical characterisation

Supplementary Movie 18: Representative video of a day 60 chronic EPS-NMO (stable) during electrical stimulation for biomechanical characterisation

Supplementary Movie 19: Representative video of a day 60 chronic EPS-NMO (increasing) during electrical stimulation for biomechanical characterisation

## Supplementary Tables

**Supplementary Table 1.**
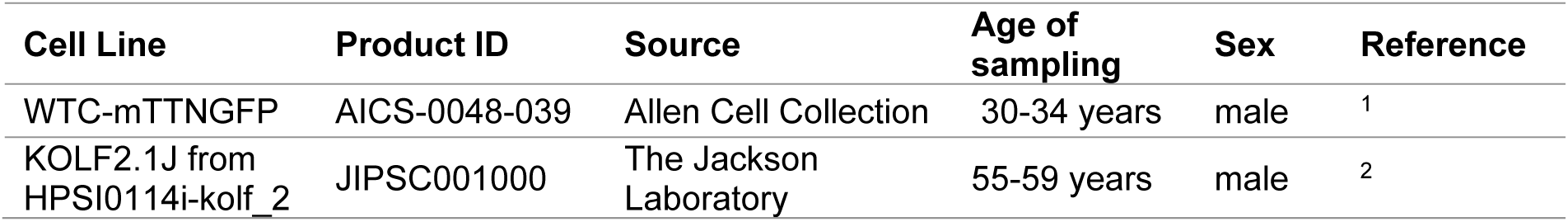
hiPSC lines.

| Cell Line | Product ID | Source | Age of sampling | Sex | Reference |
| --- | --- | --- | --- | --- | --- |
| WTC-mTTNGFP | AICS-0048-039 | Allen Cell Collection | 30-34 years | male | <sup>1</sup> |
| KOLF2.1J from HPSI0114i-kolf_2 | JIPSC001000 | The Jackson Laboratory | 55-59 years | male | <sup>2</sup> |

**Supplementary Table 2.** Primary and secondary antibodies for immunofluorescence staining.

| Antibody | Manufacturer/Source | Identifier | Dilution |
| --- | --- | --- | --- |
| Goat anti-choline acetyltransferase (ChAT) | Millipore | AB144P | 1:200 |
| Rabbit anti-cleaved Caspase-3 | Cell Signaling | 9661S | 1:1000 |
| Mouse anti-neurofilament H (SMI32) | Biolegend | 801701 | 1:1000 |
| Mouse anti-Myosin Skeletal Fast (Fast Myosin Heavy Chain; MYH1/MYH2) | Sigma Aldrich | M4276 | 1:500 |
| Rabbit anti-beta Tubulin 3/ Tuj1 (TUBB3) | Biozol | GTX129913 | 1:1000 |
| Mouse anti-Myogenic Differentiation 1 (MyoD1) | BD | 554130 | 1:50 |
| Mouse anti-Paired Box 7 (Pax7) | DSHB | PAX7-s | 1:50 |
| Goat anti-Sox1 | R&D | AF3369 | 1:500 |
| Rabitt anti-Ki67 | abcam | ab15580 | 1:501 |
| Rabbit anti-S100b | abcam | ab52642 | 1:500 |
| Rabitt anti-GFAP | Sigma Aldrich | SAB5600060 | 1:500 |
| Alexa 647 Conjugate $\alpha$ bungarotoxin | Thermo Fisher | B35450 | 1:1000 |
| Donkey anti-mouse, Alexa Fluor 488 | Invitrogen | A21201 | 1:500 |
| Donkey anti-rabbit, Alexa Fluor 568 | Invitrogen | A10042 | 1:500 |
| Donkey anti-mouse, Alexa Fluor 568 | Invitrogen | A10037 | 1:500 |
| Donkey anti-mouse Alexa Fluor 647 | Invitrogen | A31571 | 1:500 |
| Donkey anti-goat, Alexa Fluor 647 | Invitrogen | A21447 | 1:500 |

**Supplementary Table 3.** Software used in bulk RNA sequencing data analysis.

| Software (version) | Citation |
| --- | --- |
| FastQC (v0.12.1) | <sup>3</sup> |
| MultiQC (v1.25.1) | <sup>4</sup> |
| HISAT2 (v2.2.1) | <sup>5</sup> |
| samtools (v1.22.1) | <sup>6</sup> |
| featureCounts (Subread v1.5.3-0) | <sup>7</sup> |
| R version 4.4.3 (2025-02-28) | R Core Team (2024). R: A Language and Environment for Statistical Computing. R Foundation for Statistical Computing, Vienna, Austria (https://www.R-project.org/). |
| RStudio 2024.12.1+563 "Kousa Dogwood" Release | Posit team (2024). RStudio: Integrated Development Environment for R. Posit Software, PBC, Boston, MA. URL http://www.posit.co/. |
| DESeq2 (v1.46.0) | <sup>8</sup> |
| fgsea (v1.32.4) | <sup>9</sup> |
| enrichR (v3.4) | Jawaid W (2025). enrichR: Provides an R Interface to 'Enrichr'. R package version 3.4, (https://CRAN.R-project.org/package=enrichR). |
| **Reference file** | **Link** |
| Human GRCh38<br>(GENCODE<br>v49/Ensembl115<br>annotations) | https://ftp.ensembl.org/pub/release-115/fasta/homo_sapiens/dna/ |
| Homo sapiens GTF<br>(Ensembl release-115 ) | https://ftp.ensembl.org/pub/release-115/gtf/homo_sapiens/Homo_sapiens.GRCh38.115.gtf.gz |

**Supplementary Table 4.** Reagents and consumables.

| **Chemicals, Small molecules, reagents** | **Manufacturer/Source** | **Identifier** |
| --- | --- | --- |
| 2-Mercaptoethanol | Gibco, Thermo Fisher Scientific | 31350010 |
| Accutase | Sigma-Aldrich | A6964 |
| B27 | Gibco, Thermo Fisher Scientific | 17504044 |
| BAMBANKER | Nippon | BB05 |
| BSA | VWR | A7906 |
| BSA for molecular biology | Sigma-Aldrich | A9418 |
| CHIR 99021 | Tocris | 4423/10 |
| Curare | Sigma-Aldrich | T2379 |
| DAPI | Thermo Fisher Scientific | D1306 |
| Direct-zol RNA Miniprep Plus Kit | ZYMO RESEARCH | R2071 |
| DMEM F12 | Gibco, Thermo Fisher Scientific | 21331046 |
| DPBS | Gibco, Thermo Fisher Scientific | 14190-169 |
| bFGF | in house |  |
| Geltrex LDEV-Free, hESC-Qualified | Gibco, Thermo Fisher Scientific | A1413302 |
| HGF | Peprtech | 100-39H |
| IGF | Peprtech | 100-11 |
| Matrigel hESCQualified Matrix | Gibco, Thermo Fisher Scientific | 354277 |
| mTeSR | Stemcell Technologies | 85850 |
| N2 | Gibco, Thermo Fisher Scientific | 17502048 |
| NB | Gibco, Thermo Fisher Scientific | 21103049 |
| Paraformaldehyde | VWR | ALFAJ19943.K2 |
| Pen / Strep | Gibco, Thermo Fisher Scientific | 15140122 |
| Triton-X 100 | Sigma-Aldrich | T9284 |
| Trypan Blue | Gibco, Thermo Fisher Scientific | T10282 |
| Y-27632 (hydrochloride) | BioMol GmbH | Cay10005583-50 |
| Gelatin | Sigma-Aldrich | G1890 |
| Sucrose | Sigma-Aldrich | 84097 |
| Immu-Mount mounting medium | erpedia | 9990402 |
| 50 ml Falcon tubes | NeoLab | 352070 |
| 15 ml Centrifugation tubes | FAUST | TPP91015 |
| 60mm cell culture dishes | Sigma-Aldrich | CLS430166-500EA |
| 96-well U-bottom ultra-low attachment plates | Thermo Fisher Scientific | 174929 |
| Countess™ chamber | Thermo Fisher Scientific | C10228 |
| Cryovials | Thermo Fisher Scientific | 10418571 |
| Falcon® 6 Well TC-Treated Multiwell Cell Culture Plate | NeoLAB | 353046 |
| Microtome blades Type C35 | pfm medical | 207500003 |
| Superfrost Plus adhesion microscope slides | Epredia | J1800AMNZ |
| Cover slips, rectangular, 0.13-0.16mm thick | Th. Geyer | 7695031 |
| LoBind tubes 1.5 mL | VWR | 525-0133 |

**Supplementary Table 5.** Equipment and software.

| Software | Manufacturer/Source |
| --- | --- |
| Microsoft Word 16.99.2 | Microsoft |
| Micosoft Excel 16.99.2 | Microsoft |
| GraphPad Prism 9 and 10 | GraphPad Software |
| Fiji Image J 2.3.0/1.53q, 1.54p | ImageJ |
| miniforge3, conda version: 24.11.3 | conda-forge |
| Python 3.9.21 | Python.org |
| Ilastik 1.4.0.post1 | Prof. Fred Hamprecht, University of Heidelberg, ilastik.org |
| Harmony 5.2 | Perkin Elmer |
| BioRender | BioRender.com |
| Leica Application Suite (LAS) 3.5.7.23225 | LEICA |
| Enersight (v1.1.0.560) | Leica Microsystems CMS GmbH |
| PyCharm 2022.2.2 | Python.org |
| Mendeley Reference Manager (v2.138.0) | mendeley.com |
| **Equipment** | **Manufacturer/Source** |
| Countess™ | Thermo Fisher Scientific |
| Leica SP8 Confocal microscope | LEICA |
| IonOptix C-Pace EM | IonOptix/CytoCypher B.V. (Europe) |
| Opera Phenix Plus | PerkinElmer, Revvity |
| Orbital Shaker | Edmund Bühler GmbH |
| NanoDrop 2000C | Thermo Fisher Scientific |
| NovaSeq X Plus system | Illumina |
| Centrifuge |  |
| Leica DMI1 Stand, with FLEXACAM C1 | Leica Microsystems CMS GmbH |

